# Nitrosation of CD36 regulates endothelial function and serum lipids

**DOI:** 10.1101/2024.04.09.588733

**Authors:** Melissa A. Luse, Wyatt J. Schug, Luke S. Dunaway, Shruthi Nyshadham, Skylar A. Loeb, Alicia Carvalho, Rachel Tessema, Caitlin Pavelic, T.C. Stevenson Keller, Xiaohong Shu, Claire A. Ruddiman, Anna Kosmach, Timothy M. Sveeggen, Ray Mitchell, Pooneh Bagher, Richard D. Minshall, Norbert Leitnger, Linda Columbus, Kandice R. Levental, Ilya Levental, Miriam Cortese-Krott, Brant E. Isakson

**Affiliations:** Robert M. Berne Cardiovascular Research Center, University of Virginia School of Medicine; Department of Molecular Physiology and Biophysics, University of Virginia School of Medicine; Department of Pharmacology, University of Virginia School of Medicine, Charlottesville, VA, USA; College of Pharmacy, Dalian Medical University, Dalian China; Department of Cellular and Integrative Physiology, University of Nebraska Medical Center, Omaha, Nebraska, USA; Department of Pharmacology, University of Illinois-Chicago; Department of Chemistry, University of Virginia; Heinrich Heine University, Dusseldorf, Germany

## Abstract

During obesity, endothelial cells (ECs) become lipid laden leading to endothelial dysfunction. We demonstrate endothelium downregulates caveolin-1 (Cav1) in mouse and human in response to obesity. Using an EC-specific Cav1 knockout mouse, we find mice are hyperlipidemic regardless of diet, but retain endothelial cell function. Whereas initially this was thought to be due to Cav1 mediate endocytosis, we find instead the mice have significantly increased nitric oxide (NO) in response to the lack of Cav1. The presence or absence of NO toggled inversely EC lipid content and plasma lipid in mice. We found the fatty acid translocase CD36 was directly nitrosated by endogenous NO at the same cysteines that are palmitoylated on CD36. The nitrosation of CD36 prevented it’s trafficking to the plasma membrane and decreased lipid uptake. The physiological effect of this mechanism was a reliance on NO for endothelial function. This work suggests that CD36 nitrosation occurs as a protective mechanism to prevent EC lipotoxicity and preserve function.

**Teaser:** Nitric oxide regulates serum lipids and endothelial cell lipid content through nitrosation of CD36.

## Introduction

Endothelial cells (ECs), strategically positioned as gatekeepers of peripheral tissues, play a crucial role in the uptake and transportation of materials from the bloodstream to tissues^1–3^. In the context of metabolic syndrome, characterized by a state of nutritional excess, the finely tuned transport system of ECs faces a challenge. Excessive fatty acid cargo transported into the endothelium can disrupt the equilibrium of fatty acid storage and handling, leading to endothelial dysfunction^4, 5^. As a consequence, lipids begin to accumulate within the endothelium, instigating a state of lipotoxicity that further contributes to and exacerbates endothelial dysfunction^6–8^. In metabolic syndrome, the intricate process of how ECs normally take in lipids becomes dysregulated, emphasizing the importance of understanding these mechanisms to inhibit the onset and progression of endothelial dysfunction^9^. ECs possess inherent protective mechanisms that aim to combat lipotoxicity and dysfunction; however, these compensatory mechanisms fail as the lipid burden increases. This lipid-dependent dysregulation not only compromises the physiological functions of ECs but also sets the stage for the development of various cardiovascular diseases associated with obesity^10, 11^.

Caveolae, specialized membrane invaginations have long been implicated in cardiovascular and metabolic syndrome, with historical relevance attributed to their role in low-density lipoprotein (LDL) transcytosis, promoting atherogenesis^12–16^. Caveolin-1 (Cav1) emerges as a central figure in the intricate landscape of EC function, particularly through its association with caveolae. Recent investigations have unveiled additional layers of complexity in Cav1’s regulatory functions, demonstrating its role in both lipid uptake and lipolysis^4, 17–19^. Notably, Cav1 extends its influence beyond lipid metabolism and transport within ECs to include the deliberate regulation of endothelial nitric oxide synthase (eNOS)^20, 21^. Cav1 has been recognized as an integral modulator of eNOS activity and the subsequent production of nitric oxide (NO), a critical mediator of vascular homeostasis^22, 23^. The absence of caveolae disrupts the normal inhibitory relationship between Cav1 and eNOS, leading to an over production of NO. This negative regulation can be modified to aptly tune NO concentrations in ECs. Here, we draw a link between the lipid handling and eNOS regulatory roles of Cav1 establishing, for the first time, a role for NO in lipid handling. This new role for NO emphasizes the coordinated interplay between Cav1, NO, lipid uptake, and EC dysfunction.

Lipid uptake into non-fenestrated endothelium requires the transfer of fatty acids from the circulation into ECs^24^ and regulation by key proteins, including lipoprotein lipase, scavenger receptor-B1, and notably, CD36^25^. CD36 is abundantly expressed in the endothelium and serves as a crucial fatty acid transporter, allowing for the uptake of lipids into ECs^24, 26, 27^. In humans, polymorphisms in the CD36 gene are associated with hyperlipidemia, metabolic syndrome, and type 2 diabetes, highlighting the clinical relevance of CD36 in metabolic homeostasis^28^. However, the exact regulatory mechanisms behind its function within the endothelium remain not fully understood. One proposed mechanism for CD36 regulation is through the posttranslational modification palmitoylation^29–31^. Palmitoylation is a reversible modification which plays an important role in shuttling modified proteins to and from the plasma membrane. The reversible cycles of palmitoylation and depalmitoylation allow for dynamic re-localization of proteins within the cell^32–35^. Specifically, work by Hao et al. show APT1, acyl-protein thioesterase, to cleave palmitoyl groups from CD36 driving the internalization of CD36 from the plasma membrane^29^.

Proposed here, is novel post translational modification of CD36, nitrosation. Nitrosation is an important post translational modification which can alter protein localization and function^36–38^. Like palmitoylation, nitrosation is readily reversible^39^ allowing for rapid protein modifications to occur altering cellular function. Here, we present nitrosation of CD36 via endogenous NO by loss of Cav1 inhibition in obesity. We find this post translational modification works in tandem with palmitoylation as a competitive regulatory mechanism for controlling lipid transport in the adipose endothelium. Our evidence suggests the cellular changes in CD36 post translational modifications between palmitoylation and nitrosation in endothelium regulate blood lipids.

## Results

### Caveolin 1 expression decreases in endothelial cells during metabolic syndrome in mouse and human

We initially sought to identify EC transcripts that may be involved with lipid transport into endothelium during obese conditions. Using our scRNAseq of normal chow (NC) and high fat diet (HFD) adipose^40^, we subsetted the isolated ECs based on expression of specific marker genes known to be present in endothelium (**Supplementary figure 1A-F**). ECs were then further classified by their vascular origin (i.e. arterial, venous, capillary, lymphatic; **Supplementary figure 1B**). Differential gene expression analysis was performed and a reduction in Cav1 expression in HFD mice (**Figure 1A**) was observed in ECs from all vascular clusters (**Figures 1B-C**). We next used a published scRNAseq data set of human adipose tissue from both healthy individuals and patients with metabolic syndrome^41^ and also observed *CAV1* expression was decreased in adipose ECs from obese patients (**Figure 1D-F**). The diet dependent manner of this gene expression data suggested Cav1 may be an important component to lipid handling during obesity.

**Figure 1:**
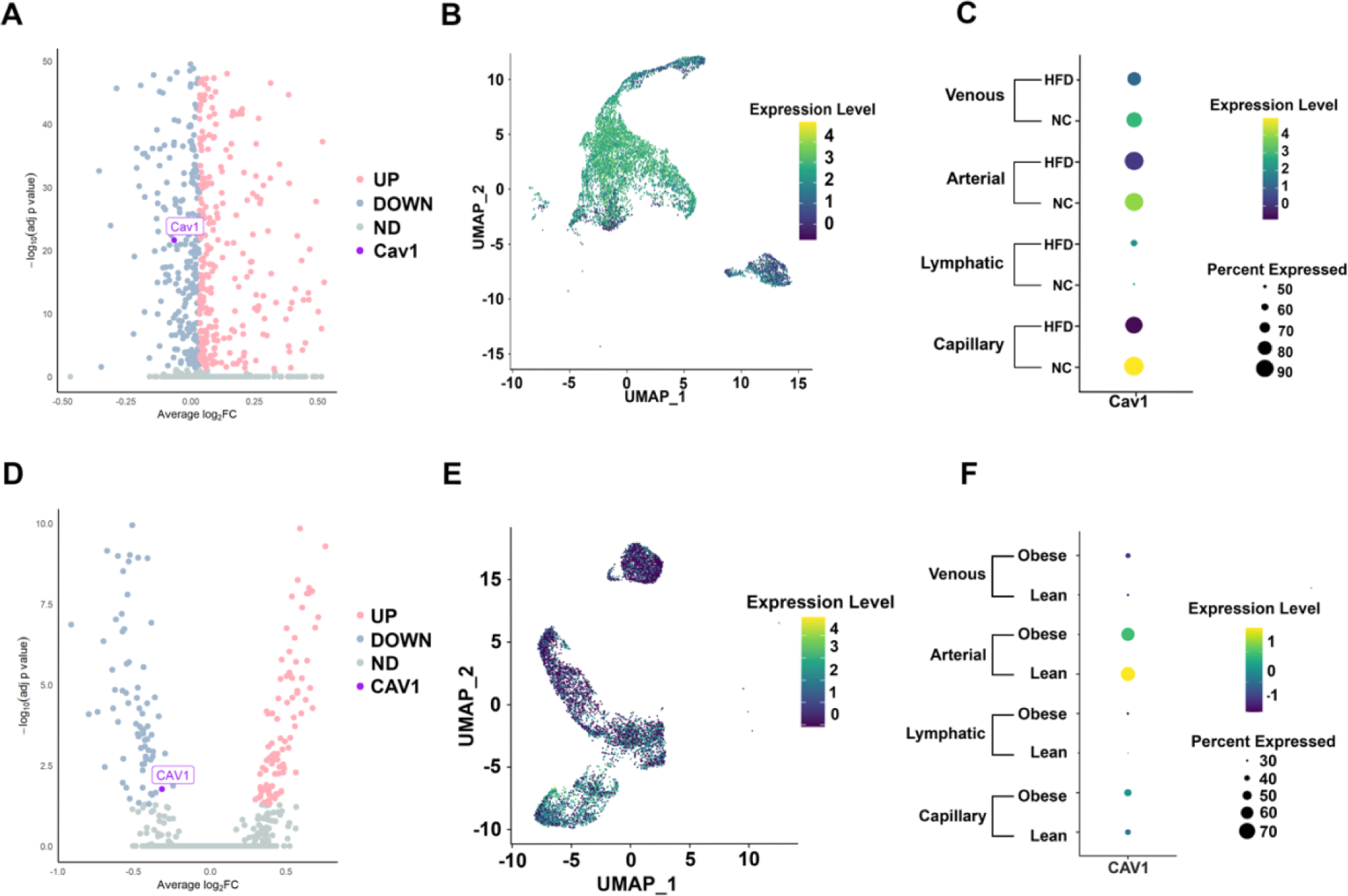
Adipose endothelial cell caveolin 1 decreases with metabolic disease. (**A**) Volcano plot showing differential gene expression in adipose ECs from HFD-fed mice relative to NC-fed mice fed over 12 weeks; genes are displayed as up-(red) or down-regulated (blue). (**B**) UMAP of murine EC *Cav1* expression across vascular clusters. (**C**) Dot plot showing *Cav1* expression among ECs of different vascular origins and diet-fed mice. (**D**) Volcano plot showing differential gene expression in human adipose ECs from obese individuals relative to lean individuals; genes are displayed as up-(red) or down-regulated (blue). (**E**) UMAP of human EC *CAV1* expression across vascular clusters. (**F**) Dot plot showing human *CAV1* expression among ECs of different vascular origins from both lean and obese humans.

### Endothelial cell Caveolin 1 can regulates serum and cellular lipids

To investigate how the down regulation of Cav1 contributes to metabolic syndrome progression, we selectively deleted Cav1 from endothelium (**Figure 2A**). Loss of Cav1 in ECs was extensively validated (**Figure 2B-D**; **Supplementary figures 2A-B**). Importantly, Cav1 protein levels in adipocytes were unaltered (**Supplementary figure 2C**). Deletion of Cav1 from endothelium did not change mouse weight, epididymal fat pad mass or food consumption (**Supplementary figures 2D-G**). Despite the lack of weight gain phenotype, EC Cre^+^ *Cav1^fl/fl^* mice when compared to EC Cre^-^ *Cav1^fl/fl^*mice, had significantly increased serum triglycerides, cholesterol, and LDL regardless of diet (**Figures 2E-G**). Importantly, EC Cre^+^ *Cav1^fl/fl^*mice fed a NC diet became hyperlipidemic, even surpassing triglyceride levels of HFD controls. HDL levels were unchanged and non-esterified fatty acids (NEFA) levels were unaffected by loss of Cav1 from ECs (**Figures 2H-I**). Next, adipose arteries were stained with Nile red to visualize en face intracellular lipid accumulation in endothelium. Arteries from EC Cre^+^ *Cav1^fl/fl^* mice had significantly decreased levels of lipid droplets relative to EC Cre^-^ *Cav1^fl/fl^* controls (**Figures 2J**). Thus, it appears the lack of Cav1 in endothelium prevents lipid accumulation in endothelial cells and promotes the accumulation of lipids in serum (**Figure 2K)**. Despite the hyperlipidemic phenotype of the mice, their ability to process glucose was improved regardless of diet (**Figure 2L, Supplementary figure 2H**). The enhanced glucose sensitivity provided initial evidence the endothelium remained more metabolically healthy compared to their EC Cre^-^ *Cav1^fl/fl^* littermates, possibly due to decreased intracellular lipid accumulation from lack of Cav1.

**Figure 2:**
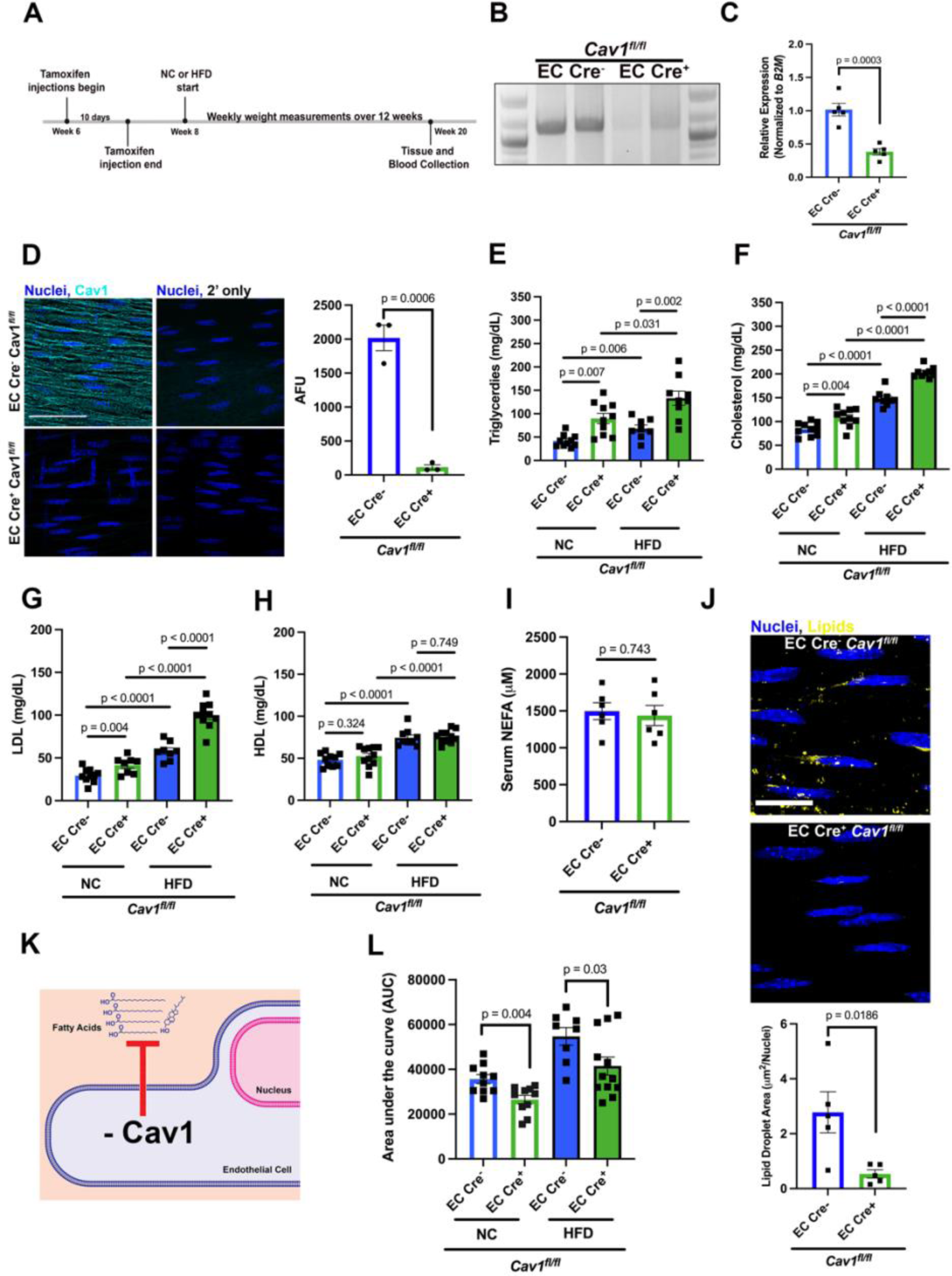
Genetic deletion of caveolin 1 in endothelial cells increases serum lipids and decreases intracellular lipids. (**A**) Schematic outlining the timeline and diet schedule for *Cav1^fl/fl^* mice. (**B**) gDNA excision gel of the Cav1 second exon from lung. (**C**) qPCR of Cav1 RNA isolated from lung. (**D**) *en face* immunostaining of adipose arteries. N= 3 (each averaged from 3 fields of view), Scale bar = 20µm, (right) quantification in arbitrary fluorescence units (AFU). (**E**) triglycerides, (**F**) cholesterol, (**G**) LDL, (**H**) HDL, (**I**) fasting non-esterified fatty acids (NEFA) from serum of EC Cre^-^ *Cav1^fl/fl^* and EC Cre^+^ *Cav1^fl/fl^* fed a normal chow (NC), N= 8-10 mice per group. (**J**) Lipid staining (Nile red) in an *en face* adipose artery quantified (below) using lipid droplet area per nuclei. N=5 mice (average from 3 fields of view each), Scale bar = 10 µm. (**K**) Diagram showing loss of Cav1 inhibits lipid uptake. (**L**) Glucose tolerance test (GTT) area under the curve (AUC) quantification. N = 8-10 mice. P-values listed are from unpaired t-tests.

We next examined the mechanism underlying the observed reduction in endothelial lipid uptake with human adipose microvascular endothelial cells (HAMECs). We recapitulated decreased lipid uptake in HAMECs with Cav1 deficiency (**Figure 3A-C**). Because caveolin organizes cholesterol in the plasma membrane into distinct lipid domains^42^, we hypothesized that the disruption of the cholesterol would also alter lipid uptake. To test this, we treated HAMECs with dimethyl-β-cyclodextrin (Cyclo) and observed similar results to the depletion of Cav1 (**Figure 3D-F**). Because lack of Cav1 and disruption of lipid domains inhibited lipid uptake, we predicted that endocytosis inhibitors would also inhibit lipid uptake. However, using two different endocytosis inhibitors (dynosore and genistein), we observed no change in lipid uptake after treatment with lipids (**Figure 3G-H**).

**Figure 3:**
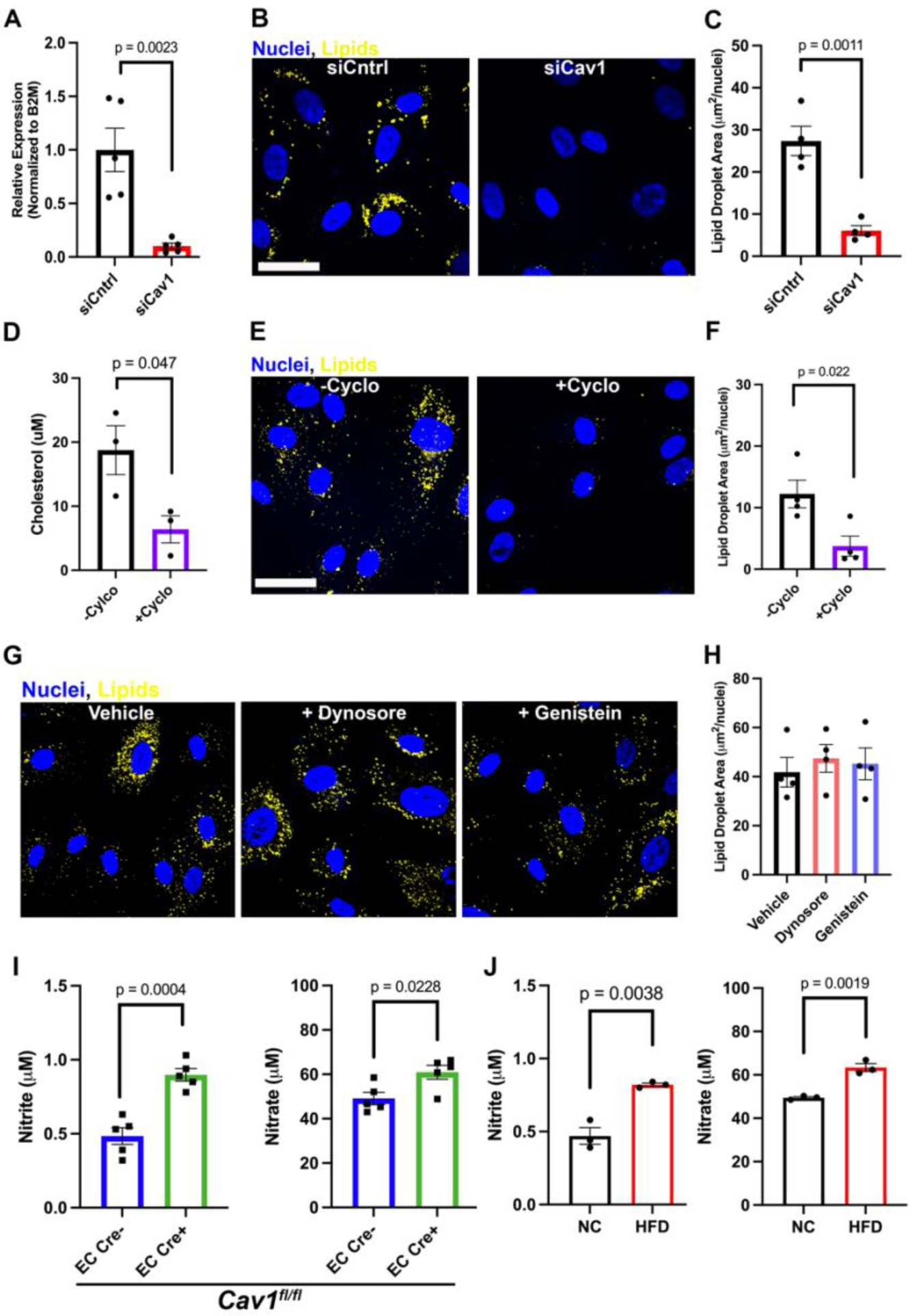
Potential role for nitric oxide, but not endocytosis, in the absence of endothelial cell caveolin 1. (**A**) Knockdown of Cav1 RNA shown via qPCR. (**B**) Lipid (Bodipy) staining in human adipose microvascular endothelial cells (HAMECs), quantified in (**C**), N= 4 (with each as an average of 3 fields of view). (**D**) Cholesterol concentration after cyclodextrin treatment (+cylco, 10mM) or control treatment (-cyclo). (**E**) Lipid staining (Bodipy) of HAMECs after cyclodextrin and lipid treatments, quantified in (**F**). (**G**) Lipid (Bodipy) staining and quantification (**H**) of cells treated with vehicle or endocytosis inhibitors (dynosore (80µM) and genistein (200µM)) before lipid treatment. (**I**). Nitrite (left) and nitrate (right) levels in EC Cre^-^ *Cav1^fl/fl^*and EC Cre^+^ *Cav1^fl/fl^* mice, N= 5 mice. (**J**) Nitrite and nitrate levels in NC- and HFD-fed mice at 12 weeks of diet. N = 3 mice per group. 50µM lipids were used for treatments in all cells for 15 minutes. Scale bars = 25µm throughout. P-values listed are from unpaired t-tests. 50µM lipids (12:5µM linoleic acid, 25µM oleic acid, 12.5µM palmitic acid)

### A potential role for endothelial nitric oxide in regulating lipids

It is well described there is an increase in nitric oxide (NO) in the absence of Cav1 due to its role in eNOS inhibition^20, 21, 43–45^. An increase in NO is also observed with the depletion of cholesterol in the plasma membrane which disrupts Cav1-eNOS interactions.^46–48^ At 20-weeks of age (12 weeks on NC diet) we measured nitrite and nitrate levels (nitrite and nitrate are stable oxidation products of NO) from *Cav1^fl/fl^* mice and found EC Cre^+^ *Cav1^fl/fl^* mice had significantly increased serum nitrite and nitrate (**Figure 3I**). Furthermore, mice on a HFD that we demonstrated in Figure 1 had a loss of Cav1 in adipose EC, also had significantly increased nitrite and nitrate (**Figure 3J**). These results suggested NO derived from loss of Cav1 could regulate endothelial lipid uptake.

To test the hypothesis the Cav1 loss is causing a compensatory increase in NO and disrupting lipid uptake, we used HAMECs genetically devoid of Cav1, a condition that previously inhibited lipid uptake (Figure 3). We treated these cells with L-NAME, a well-known inhibitor of nitric oxide synthase^49, 50^, to examine the role of NO in endothelial lipid uptake. After lipid treatment, cells devoid of Cav1 were indistinguishable from control treated cells (**Figure 4A**). We repeated this experiment *in vivo* by adding L-NAME in the drinking water of EC *Cav1^fl/fl^*mice for 2-weeks. Adipose arteries from EC Cre^+^ *Cav1^fl/fl^* mice had lipid droplets present in the endothelium (**Figure 4B**), a result strikingly different than the EC Cre^+^ *Cav1^fl/fl^* that did not receive L-NAME water (Figure 2). These data indicate decreased lipid uptake from the loss of EC Cav1 may be rescued with the inhibition of nitric oxide production.

**Figure 4:**
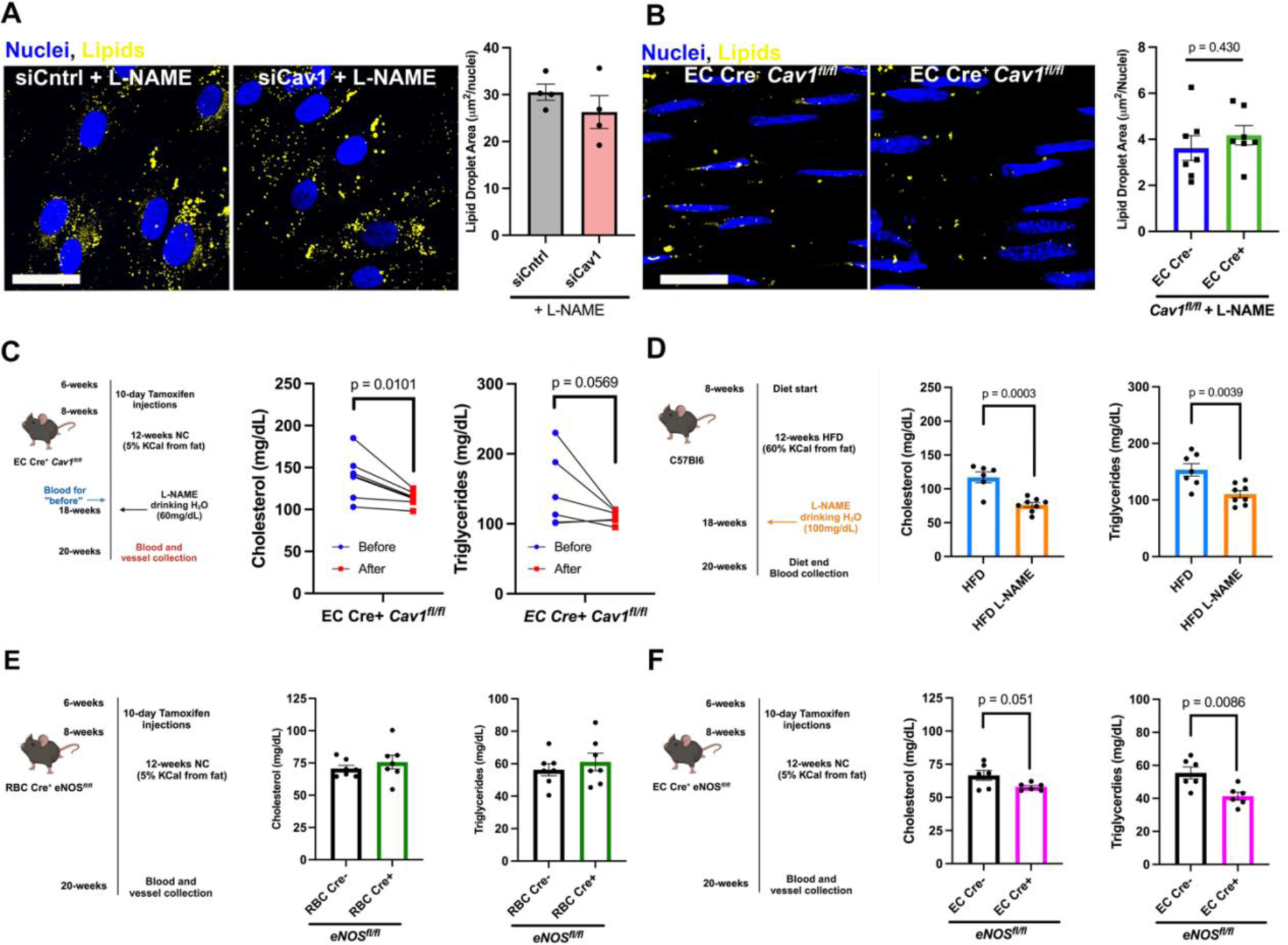
L-NAME rescues loss of lipid uptake seen with deletion of endothelial caveolin 1. (A) HAMECs treated with control (siCntrl) or Cav1 (siCav1) siRNA followed by L-NAME and stained for lipid accumulation (Bodipy) after treatment with 50µM lipids (50µM lipids (12:5µM linoleic acid, 25µM oleic acid, 12.5µM palmitic acid). N=4 (each from an average of 3 fields of view). (**B**) Lipid staining (Nile red) in *en face* adipose arteries of EC Cre^-^ *Cav1^fl/fl^* and EC Cre^+^ *Cav1^fl/fl^* mice given L-NAME (60mg/dL) in their drinking water. N=7 (each from an average of 3 fields of view). (**C**) Cholesterol and triglyceride concentrations from blood taken before and after EC Cre^+^ *Cav1^fl/fl^* mice were administered L-NAME in their drinking water. N=5-6. P-values listed are from paired t-tests. (**D**) Cholesterol and triglyceride concentrations from mice fed a high fat diet (HFD) and administered either regular drinking water or L-NAME water. (**E**) Cholesterol and triglyceride concentrations from RBC Cre^-^ *eNOS^fl/fl^* and RBC Cre^+^ *eNOS^fl/fl^* mice. N=6-7. (**F**) Cholesterol and triglyceride concentrations from EC Cre^-^ *eNOS^fl/fl^* and EC Cre^+^ *eNOS^fl/fl^* mice. N=6. P-values listen for (**D-F**) are from unpaired t-tests.

Since the EC Cre^+^ *Cav1^fl/fl^* mice given L-NAME were able to accumulate lipids within the endothelium, we hypothesized that their blood lipids would be decreased after drinking L-NAME water relative to before the onset of treatment. We took blood before giving the mice L-NAME water and then took blood after their 2-week treatment and saw a significant decrease in both serum triglycerides and cholesterol after L-NAME consumption (**Figure 4C**). Similarly, HFD mice were also given L-NAME drinking water for 2-weeks and compared to HFD mice given regular water (**Figure 4D**). The L-NAME treated group had significantly lower serum triglycerides and cholesterol compared to the control water group (**Figure 4D**). To determine if the source of NO, we used *eNOS^fl/fl^* mice either on a red blood cell (RBC) or EC Cre^22^. Loss of eNOS from RBCs had no effect on serum lipids (**Figure 4E**). However, deletion of eNOS from endothelium significantly decreased serum triglycerides and lowered cholesterol (**Figure 4F**). Thus, endogenous NO from endothelium is capable of regulating serum and intracellular lipids.

### NO modification of CD36 regulates lipid uptake

Next, we sought to identify the target of NO within the endothelium that may be responsible for regulating lipid uptake. Because there was an association between changes in serum and intracellular lipid levels and changes with alterations in Cav1, we performed lipid domain fractionation and blotted for CD36, a fatty acid translocase present on endothelium^24, 27, 51^, which has been shown to colocalize with Cav1^52–54^. We also observed CD36 in caveolar enriched lipid fractions (**Figure 5A**). However, when visualized within adipose endothelial arteries en face, we find Cav1 and CD36 signals did not overlap (**Figure 5B**).

**Figure 5:**
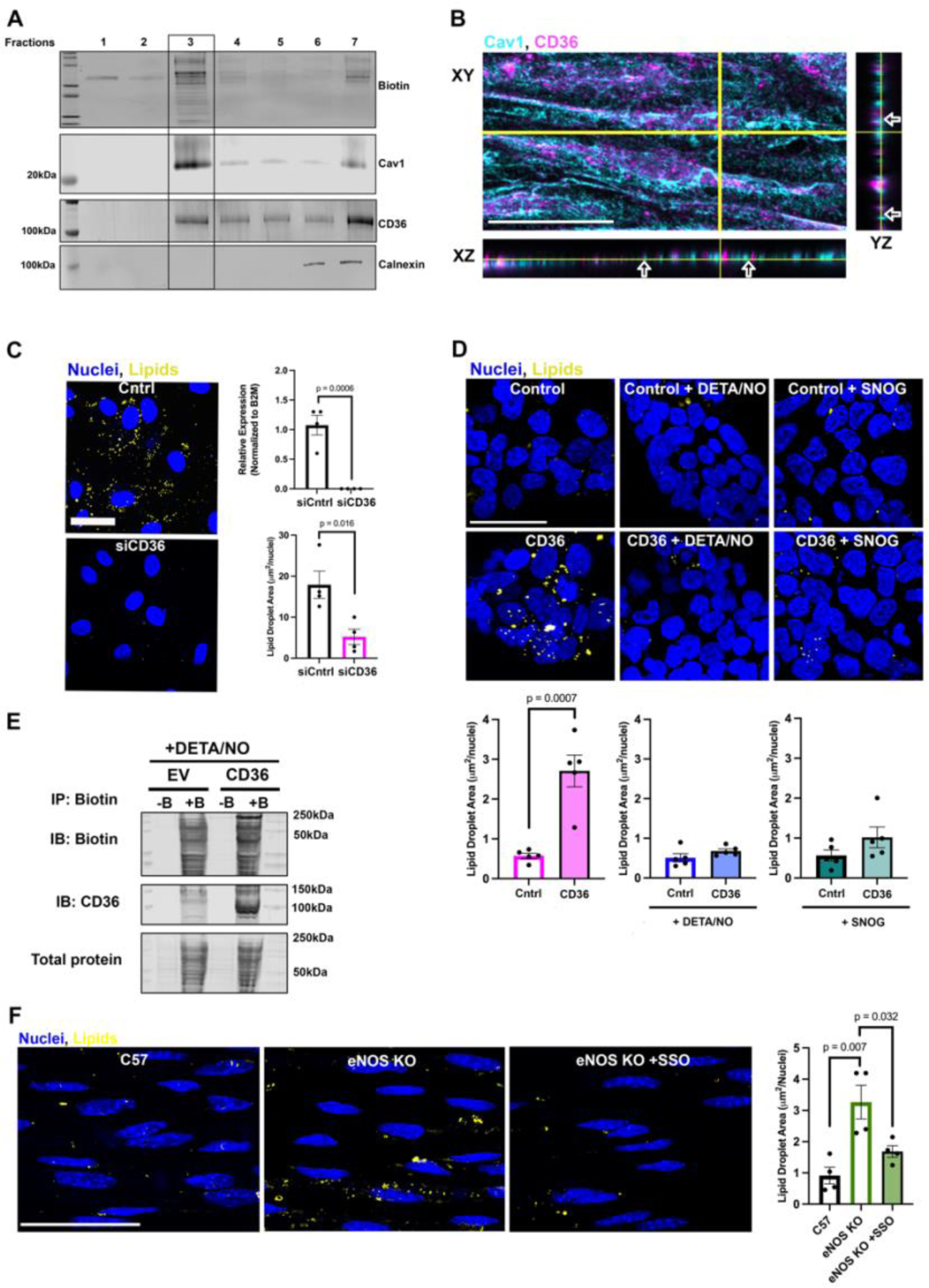
CD36 and nitric oxide may both be necessary for lipid uptake in adipose endothelial cells. (**A**) Caveolin lipid raft fractionation in HAMECs with the cell surface marked with biotin, the Cav1-rich lipid raft marked as fraction 3, and calnexin marking ER fractions. (**B**) Immunostaining of an *en face* adipose artery showing Cav1 (cyan) and CD36 (magenta) in various planes of view (XY, XZ, YZ). Arrows highlight locations in which Cav1 and CD36 are next to one another. Yellow bars show the location in which the XZ and YZ planes were taken from. Scale bar = 10µm (**C**) Control (Cntrl) vs. siCD36-treated HAMECs treated with lipids (50µM lipids; (12:5µM linoleic acid, 25µM oleic acid, 12.5µM palmitic acid)) (stained with Bodipy) and quantified. Scale bar = 25µm. (**D**) HEK cells transfected with CD36 plasmid or empty vector (control) and subsequently treated with lipids and one of two NO donors: DETA/NO or SNOG. Quantification of lipid uptake (below). Scale bar = 25µm, N=5 distinct experiments. (**E**) Biotin switch assay, with subsequent streptavidin bead pull down. All samples treated with DETA/NO. EV = empty vector, CD36 = CD36 plasmid, -B or +B = without or with biotin added, respectively. (**F**) Adipose artery *en face*, lipid accumulation quantified N = 4, three images were averaged for each value, SSO is Sulfo-N-Succinimidyl oleate a CD36 inhibitor. Scale bar = 20µm. All P-Values denoted are unpaired t-tests.

This data shows they are in lipid domains which are in close proximity to one another, but not likely within caveola (e.g.,^55, 56^). To determine if the CD36 is necessary for lipid uptake in adipose endothelium, we used CD36 siRNA mediated knockdown of CD36 (siCD36) in HAMECs. Cells treated with siCD36 had significantly decreased lipid uptake when comparted to controls (**Figure 5C**). Specificity of the CD36 antibody used are shown in **Supplementary figure 3A-B**. Cells treated with siCD36 did not have altered expression of Cav1 (**Supplementary figure 3C**). To assess whether CD36 is sufficient for lipid uptake, we used HEK293T (HEK) cells which do not endogenously express CD36 or eNOS (**Supplementary figures 3D-F**). Lipid uptake in HEK cells was only observed with CD36 transfection (**Figure 5D**). Because CD36 was sufficient for lipid uptake, and NO regulated serum and intracellular lipid accumulation independent of Cav1, we tested whether NO targeted CD36. Using an NO donor and an S-nitrosothiol (DETA/NO and SNOG), we were able to block lipid uptake in HEK cells transfected with CD36 (**Figure 5D**). A common method by which NO regulates protein function is by nitrosation (first described by Stamler et al^37, 57^). Using a biotin-switch assay^58^, we found CD36 is nitrosated in the presence of NO (**Figure 5E**). To further solidify the association between NO and CD36, we compared lipid uptake in mouse adipose arteries expressing or genetically lacking eNOS. The lack of NO from eNOS-deficient mice caused an accumulation of lipids in the endothelium (**Figure 5F**), that was blocked by the CD36 inhibitor SSO (**Figure 5F**). The sum of the data indicates increased NO, generated from lack of Cav1, may regulate CD36 lipid transport by nitrosation.

To identify which cysteines may be nitrosated on CD36, we turned to GPS-SNO^59^, a software designed to predict sites of nitrosation. We found three specific cysteines (3, 313, 466) on CD36 likely to undergo nitrosation (**Figure 6A**). These three cysteines lie in critical regions for the function and localization of the CD36 protein (**Figure 6B**). Cysteines 3 and 466 are palmitoylated^29–31^ and cysteine 313 is involved in the formation of disulfide bridges that are present in the fatty acid binding pocket of the protein^60, 61^. To determine which cysteines on CD36 may be nitrosated, we used mutated versions (C→A) of CD36 at C3, C313, and C466. First, we used a version of the plasmid where all three cysteines of interest were mutated (C466A/313A/3A; **Figure 6C**). Minimal lipid uptake was seen in cells containing the C466A/313A/3A mutant CD36. To dissect which individual or combination of cysteines was responsible for the loss of lipid uptake, individual and double mutants were made. DETA/NO treatment inhibited lipid uptake in each single cysteine mutation. We next tried a double mutation of C313A/3A and saw a decrease, however not statistically significant, in lipid uptake with DETA/NO administration. A biotin switch assay was performed, confirming that CD36 was still nitrosated in the C313A/3A CD36 mutants (**Supplementary figure 3G**). Due to the persistent nitrosation of CD36 in the C313A/3A mutants, an additional double mutation was created to probe the combinatorial role of cysteine 466 and 3 nitrosation. Mutation of C466A/3A inhibited lipid uptake mimicking the phenotype seen in the C466A/313A/3A CD36 mutant **(Figure 6C**). With the mutation of C466A/3A, nitrosation of CD36 was no longer detected via biotin switch (**Figure 6D**). The cumulative lipid uptake data (**Supplementary figure 3H**) and detection of nitrosation indicate cysteines 3 and 466 to be crucial for CD36 mediated lipid uptake.

**Figure 6:**
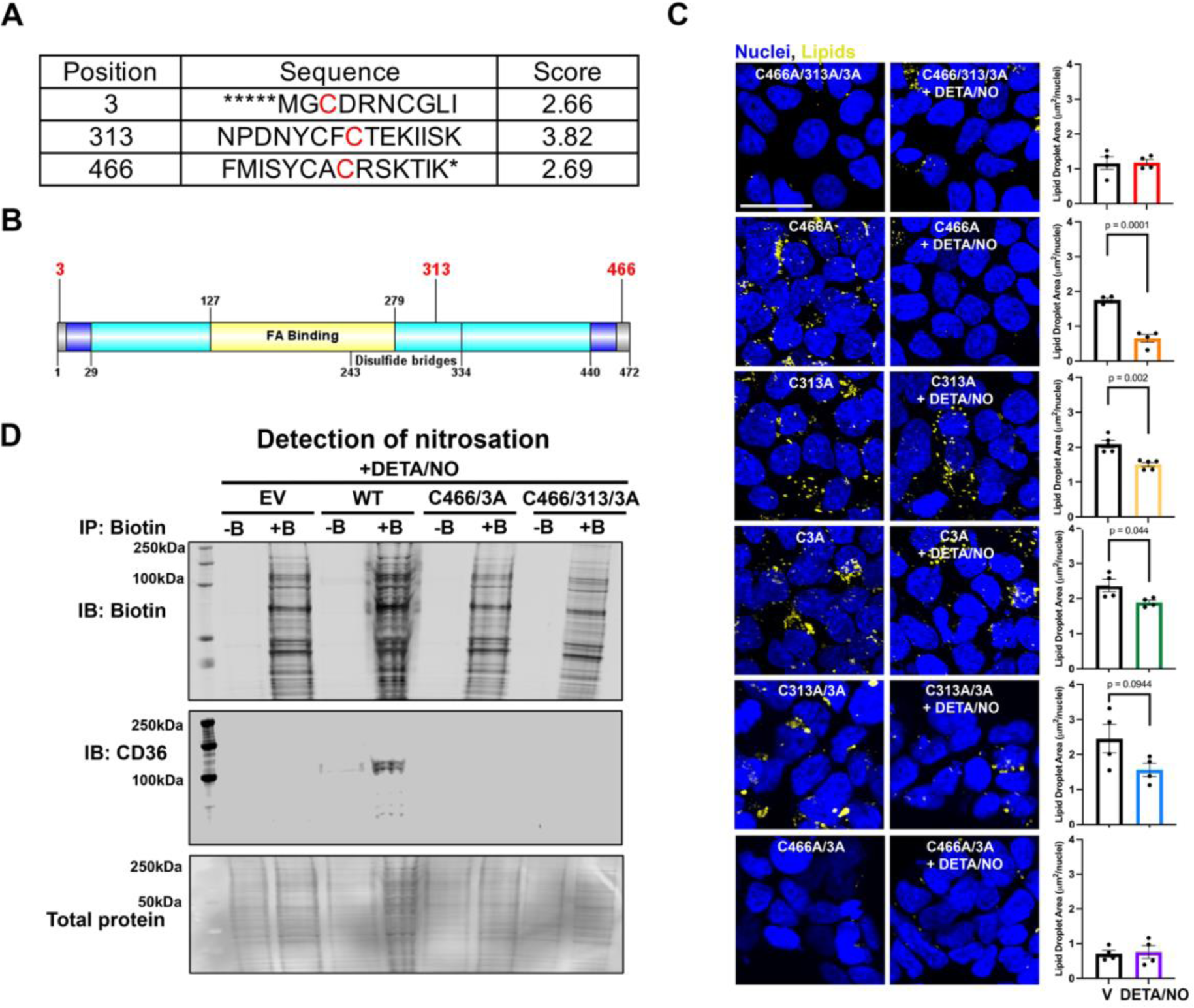
Cysteines 3 and 313 on CD36 are nitrosated inhibiting lipid uptake. (**A**) Table showing three cysteine positions on CD36 protein predicted by GPS-SNO to be potential points of nitrosation. (**B**) GPS-SNO-predicted cysteines (3, 313, 466) on CD36 protein with labeled domains. Figure created using GPS-SNO software. Yellow region denotes fatty acid binding pocket of CD36; the disulfide bridges supporting the fatty acid pocket are marked from residues 243-334. Grey denotes the intracellular domains, dark blue denotes the transmembrane regions, and teal denotes the ectodomain of CD36. (**C**) HEK cells transfected with mutated versions (C→A) of the CD36 plasmid. All three cysteine sites (3, 313, 466) on the top, then from top to bottom sites 466, 313, 3, both 313 and 3, and lastly both 466 and 3 on the bottom. Each condition was treated with 50µM DETA/NO and 50µM lipids. Quantification for each condition to the right, N = 4-5 with 3 images averaged for each N value. (**D**) Biotin switch assay, with subsequent streptavidin bead pull down. All samples treated with DETA/NO. EV = empty vector, WT = CD36 plasmid, C466/3A and C466/313/3A are mutant plasmids with mutations at specified sites, -B or +B = without or with biotin added respectively. CD36 plasmid is mCherry tagged.

Next we hypothesize that nitrosation of cysteines 3 and 466 on CD36 inhibits palmitoylation, inhibiting proper localization of CD36 to the plasma membrane and subsequently decreasing lipid uptake (**Figure 7A**). To assess CD36 localization with loss of Cav1, an environment with increased NO, we used caveolin lipid domain fractionation and siRNA for Cav1. With siCav1, CD36 is no longer enriched in the caveolin/plasma membrane fraction (**Figure 5A**: fraction 3) and is was shifted towards fraction 4 overlapping with the ER marker calnexin (**Figure 7B**). Total CD36 expression at the RNA or protein level does not change with loss of Cav1 (**Supplementary Figure 4**). If loss of Cav1 and therefore an increase in NO is disrupting the localization of CD36 this effect should be reversed with L-NAME. To test this, HAMECs were treated with either control (Cntrl) or Cav1 siRNA (siCav1) then treated with L-NAME or a vehicle control. The surface of the cells was biotinylated to capture only plasma membrane proteins. Cells lacking Cav1 had decreased plasma membrane CD36; however, cells treated with L-NAME, restored CD36 at the plasma membrane (**Figure 7C**). Because palmitoylation of CD36 is necessary for its plasma membrane localization^30^, we tested the effect of NO on palmitoylation levels with the addition of DETA/NO to HEK293T cells. Using an IP-ABE assay, we were able to detect decreased palmitoylation of CD36 in the presence of NO (**Figure 7D**). Since cysteines 3 and 466 on CD36 are important for protein localization^29^, we transfected HAMECs with either a wild type or C466A/3A mutant version of the CD36 plasmid to assess localization of the protein. HAMECs receiving the C466A/3A mutant CD36 had protein sequestration inside the cells as opposed to the plasma membrane staining seen in the wild type protein (**Figure 7E**). Additionally, cells transfected with wild type CD36 and treated with DETA/NO for 30 minutes had decreased CD36 present at the plasma membrane (**Figure 7E**). Last, we wanted to modulate CD36 localization *in vivo* using both EC Cre^-^ and EC Cre^+^ *Cav1^fl/fl^* mice and L-NAME drinking water. EC Cre^-^ mice demonstrate a diffuse and punctate staining pattern for CD36 (**Figure 7F**). Secondary only staining controls for CD36 are show in **Supplemental figure 4B**. EC Cre^+^ mice lose the punctate staining pattern and CD36 becomes compartmentally sequestered in ECs similar to a pattern of endoplasmic reticulum (ER). CD36 staining in EC Cre^+^ mice administered L-NAME have a more similar staining pattern to their EC Cre^-^ controls (**Figure 7F**) and is quantified using Pearson’s correlation coefficient. The data therefor suggests nitrosation may prevent CD36 transport to the plasma membrane and in so doing, decreases intracellular endothelial lipids, and increases plasma lipids.

**Figure 7:**
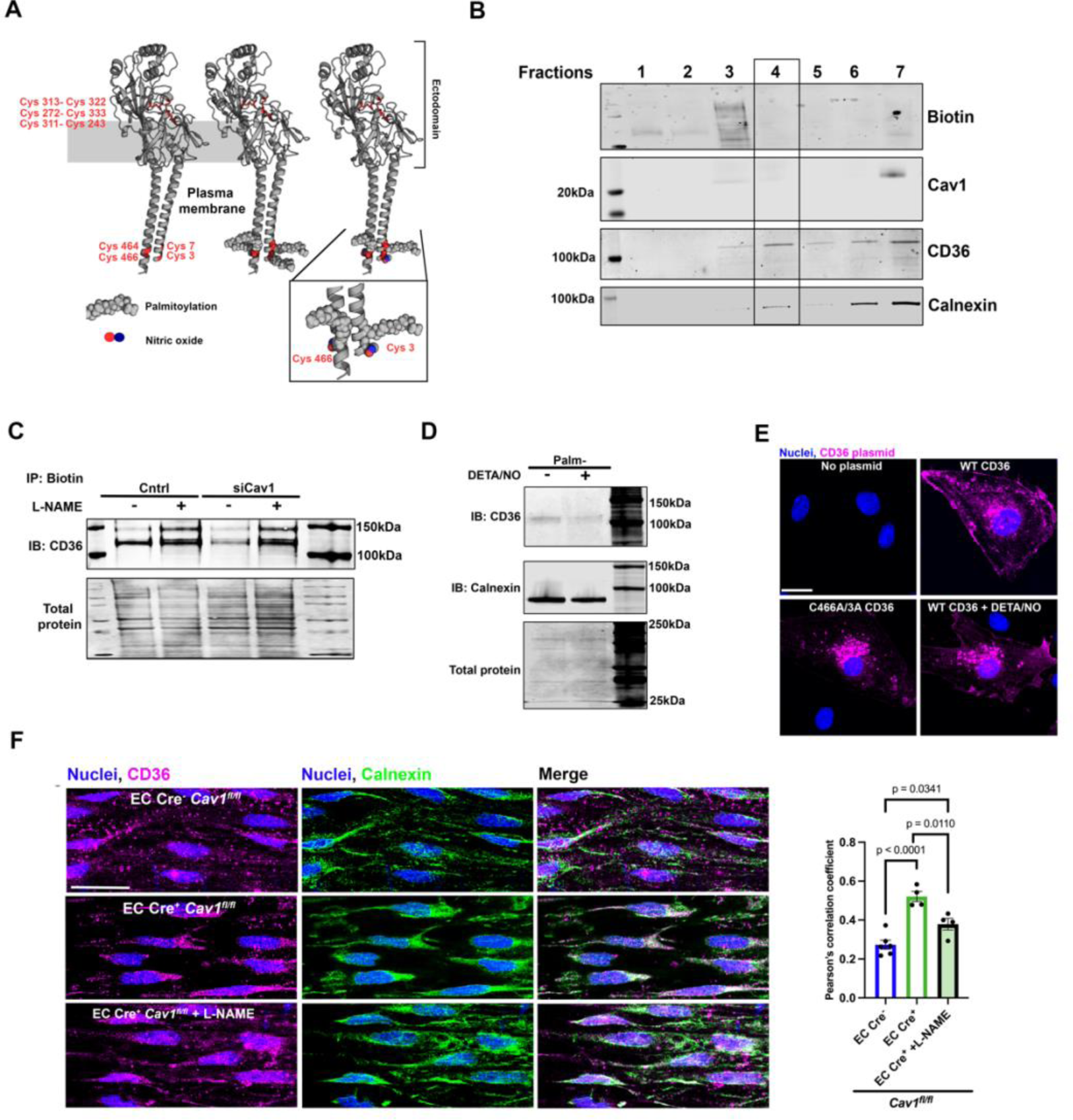
Nitrosation of CD36 inhibits its plasma membrane localization. (**A**) Structure of CD36 protein with transmembrane and ectodomains marked. Cysteines present in CD36 are marked in red; (left) cysteines in the ectodomain participate in disulfide bonds, (middle) CD36 with four cytoplasmic cysteines palmitoylated, and (right) CD36 with proposed cysteines nitrosated. (**B**) Caveolin lipid raft fractionation from HAMECs treated with Cav1 siRNA. Cav1-rich lipid rafts appear in fraction three when Cav1 is present (Figure 5A). Black box marks a shift in fractions where CD36 is enriched. The cell surface is marked with biotin and Calnexin marks ER fractions. (**C**) Membrane biotinylation with subsequent streptavidin bead pull down in HAMECs. HAMECs were treated with control siRNA (Cntrl) or siCav1 and then half were given L-NAME. (**D**) IP-ABE assay on HEK293T cells transfected with CD36-mcherry. Detection of palmitoylated proteins via western blot. (**E**) HAMECs treated with no plasmid control, WT CD36, or C466/3A mutant plasmid. DAPI denotes nuclei, and magenta marks CD36. Scale bar = 10µm. (**F**) *En face* adipose arteries from EC Cre^-^ *Cav1^fl/fl^* (top), EC Cre^+^ *Cav1^fl/fl^* (middle), and EC Cre^+^ *Cav1^fl/fl^*+ L-NAME (bottom) mice stained for CD36 (magenta) and ER with calnexin (green). Quantification of CD36-Calnexin overlap using Pearson’s correlation coefficient on the right, N= 4 mice, statistics represent one-way ANOVA with multiple comparisons.

### NOS inhibition worsens endothelial dysfunction

With an increase in EC lipid accumulation, we would expect to see endothelial dysfunction. To assess the impact of lipids on the EC transcriptome we performed bulk RNA-sequencing on HAMECs treated with lipids, L-NAME, and DETA/NO. HAMECs receiving both lipid and L-NAME treatment had a marked decrease in mitochondrial oxidative phosphorylation genes and a significant increase in lipid storage genes (**Figure 8A, Supplementary figures 5A-D**). To investigate the functional effects of decreased mitochondrial gene expression we used a Seahorse bioenergetics assay to perform a mitochondrial stress test and found HAMECs treated with lipids had a decrease in maximal oxygen consumption (**Figure 8B**), but the addition of both lipids and L-NAME decreased maximal oxygen consumption even further. Supplementing with DETA/NO, we reversed the decreased oxygen consumption and restored mitochondrial function to control levels even in the presence of lipids. This functional data supported the bulk RNA-sequencing analysis. Both loss of Cav1 and L-NAME alone did not affect mitochondrial bioenergetics (**Supplementary figures 5E-F**). To assess how this dysfunction translates to arterial function we used pressure myography to measure the vasodilatory capacity of third order mesenteric arteries. Using C57Bl/6 mice, the presence of L-NAME does not alter acetylcholine (ACh) induced vasodilation (**Figure 8C**), an effect commonly observed for small arteries^62–66^. However, if the same arteries were exposed to lipids with L-NAME, preventing nitrosation of CD36 and allowing an influx of lipids into endothelium, only then was the ACh dilation significantly decreased (**Figure 8C**). We interpret this data as evidence that NO’s effect on vascular function of small arteries may be preferentially to regulate lipid uptake via CD36.

**Figure 8:**
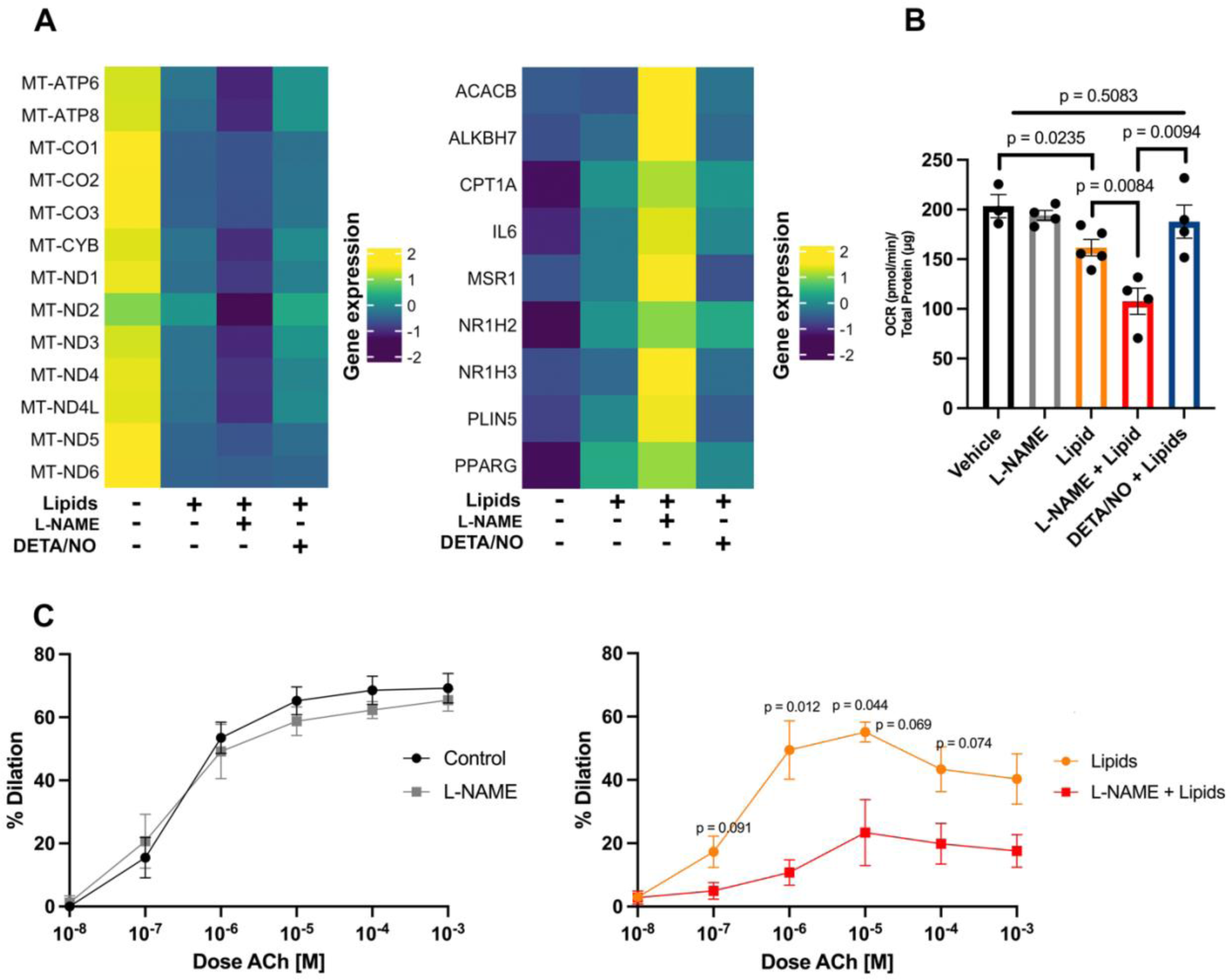
NOS inhibition drives endothelial dysfunction. (**A**) Heatmap showing enrichment of mitochondrial genes (left) in control treated cells and a decrease in expression with administration of lipids + L-NAME; statistics in supplemental figure 5A. Right, heatmap showing enrichment of lipid storage genes in L-NAME treated cells and a decrease in expression with non-lipid treated cells (control). Statistics and GSEA plot in supplemental figure 5B-E. (**B**) Maximum oxygen consumption normalized to total cellular protein from mitochondrial stress test performed via seahorse assay, N = 3-4, P values from one-way ANOVA with a multiple comparisons test. (**C**) Vasodilation of 3^rd^ order mesenteric arteries when exposed to L-NAME and/or lipids. A pre-treatment of 10µMoleic acid was used. Oleic acid was also present during treatments. N= 3-5

## Discussion

ECs are woven throughout every major organ system and facilitate the transport of critical cellular nutrients and supplies. Cellular energy needs can fluctuate, requiring ECs to constantly adapt their transport function. However, during times of excess lipid burden, such as in metabolic syndrome and obesity, this crucial energy mediator can become dysfunctional, causing other organs systems to fail. ECs must have a way to control lipid build-up or else they risk organs becoming insulin-resistant. Here, we demonstrate one mechanism in which the endothelium protects itself from excess lipid accumulation and the subsequent lipotoxicity that follows. Our data suggests a novel mechanism whereby NO modifies the fatty acid transporter CD36, preventing its localization to the plasma membrane and decreasing EC lipid accumulation (**Supplementary figure 6**). This work is a culmination of three exciting findings. First, at the cellular level, altering either palmitoylation or nitrosation of CD36 at the same cysteines regulates endothelial lipid uptake and cellular function. Second, at the tissue level, we provide evidence for the physiological importance of nitric oxide as a mediator of lipid accumulation.

Our initial data demonstrated in mouse and humans a loss of Cav1 in adipose endothelium in response to obesity. To better understand how the loss of Cav1 impacts endothelial function in obesity, we generated EC Cav1 deficient mice. Originally, we expected the phenotypes of an EC specific Cav1 deletion to be similar to the global Cav1 knockout mouse^67, 68^. Cav1 null mice are resistant to HFD-induced weight gain, insulin resistant^68^, glucose insensitive^69^, and hyperlipidemic^67^. Our EC specific Cav1 deletion had no weight gain differences compared to controls and surprisingly enhanced glucose sensitivity. Deletion of EC Cav1 and Cav1 null mice had a matching phenotype with elevated serum lipids. Many of the phenotypes associated with the Cav1 null mice have been attributed to adipocyte loss of Cav1. The data we generated helps to tease out the contributions of EC Cav1 compare to adipocyte Cav1 to whole animal metabolism.

Interestingly, the phenotypes seen with loss of EC Cav1 mimic that off an EC-specific loss of CD36. EC CD36 knockout mice have increased postprandial serum triglyceride levels, decreased intracellular lipid droplet formation, and enhanced glucose sensitivity^24^. Conversely, overexpression of CD36 in muscle reduces serum triglycerides and increases glucose insensitivity^70^. Together, this drives the idea that these two proteins exist in a regulatory pathway related to lipid entry into endothelium.

We considered two hypotheses for how Cav1 regulates lipid uptake. The first, is that Cav1 does so through endocytosis. However, we found that inhibition of endocytosis with dynosore and genistein showed no effect on lipid uptake. This finding was surprising as the literature surrounding Cav1 and lipid uptake suggests its mechanism for doing so is through endocytosis^26^. The second possibility we considered is that increased NO associated with loss of Cav1 inhibits lipid uptake. Nitrosation from NO has been shown to regulate protein function and localization^38, 71^. Therefore, we proposed that Cav1 regulates lipid uptake by modulating endothelial NO levels.

According to our results, the addition of NO to cysteines C3 and C466 on CD36 inhibits CD36 from being trafficked to the plasma membrane and instead retains the translocase within the ER. Therefore, less lipids can enter endothelium with nitrosation of CD36. Converse to nitrosation, previous work has identified palmitoyl groups as key regulators of CD36 localization^29–31, 72^. Specifically, the N and C-terminal cysteines (C3, C7, C464, C466) have been demonstrated as sites of palmitoylation for proper CD36 localization to the plasma membrane. The palmitoylated CD36 at the plasma membrane functions to take in lipids^29–31, 72^. This mechanism has been identified in HEK293T, 3T3L1, adipocytes, and macrophages^29–31, 72^. We found identical cysteines, C3 and C466, can be either nitrosated or palmitoylated on CD36. We propose ECs can use the post-translational modifications of nitrosation or palmitoylation on CD36 for localization at the plasma membrane and modulate lipid uptake.

Once lipids enter endothelium, their fate is not fully understood. Literature points to four known outcomes for lipids, re-esterification for storage in lipid droplets, direct use for ATP production through oxidative phosphorylation, remain in the cell inducing various cellular stress pathways, or transported to other tissues^9^. Here, we show robust staining of lipid droplets in endothelium both upon lipid stimulation as well as at homeostasis. We hypothesize that the excess lipids which are taken into the endothelium with NOS inhibition are both stored within lipid droplets and reside within the cytoplasm inducing cellular stress via lipotoxicity. Our mitochondrial respiration data show when lipids are added to HAMECs the cells don’t necessarily utilize the extra fatty acids for fuel. In fact, this addition of lipid decreased the cells mitochondrial function. It has been demonstrated that the alteration of Cav1 dynamics in ECs can alter mitochondrial fusion and fission dynamics directly causing mitochondrial dysfunction^73^. We hypothesize this is likely due to lipotoxicity induced by lipids that were not stored, which can ultimately affect cellular respiration. Our data highlights two destinations for lipids upon entry into endothelium.

Moving from the cell to the tissue, how does the excess lipid accumulation in EC causing mitochondrial dysfunction alter arterial dilation, the essential physiological mechanism whereby blood flow and blood pressure are regulated? Current literature focuses on NO modulating endothelial-induced vasodilation. This effect of NO has been demonstrated a plethora of times in large arteries, but in small arteries, it has become clear other mechanisms strongly predominate^62–66^. In agreement with these observations, our data shows no change in ACh induced vasodilation in the presence of L-NAME. However, it is the combined presence of lipids *and* L-NAME that ultimately decreased vasodilatory capacity. Our findings are in alignment with results from ECs treated with lipids and L-NAME, showing a dramatic decrease in mitochondrial function. Based on our data, L-NAME moves CD36 away from the plasma membrane to the ER in control and lipid treated arteries. However, it is the presence of the lipids that alter EC function by being taken up into the cells when NO, or mechanisms to make NO, is deplete. In this way, our data points to nitrosation, a crucial regulator of various cellular processes by protein post-translational modification, as a major function for NO in small arteries.^21, 23, 74^. This may also explain the difference between utilization of NO in small versus large arteries.

Currently, it is thought that elevated serum triglycerides can lead to the development of type 2 diabetes and metabolic syndrome^75, 76^. How then, do the EC *Cav1^fl/fl^*mice with elevated triglycerides have an enhanced ability to process glucose? We hypothesize that due to the decrease in intracellular lipids, ECs remain glucose sensitive and can maintain glycemic control. This would present EC lipid accumulation and lipotoxicity as an emerging predictor for the development of metabolic syndrome. This idea is well-supported by other studies which describe endothelial dysfunction contributing to the etiology of metabolic syndrome^77^. Here, we demonstrate it is the intracellular lipid accumulation in the microvasculature, rather than elevated serum triglycerides, that plays a larger role in the onset of endothelial dysfunction.

Our work is not without alternative explanations. For example, the increased NO metabolites measured in our study could be a byproduct of peroxynitrite activity^78^. Vascular peroxynitrite has been shown to increase with obesity^79, 80^. Additionally, loss of Cav1 under pathological conditions has been linked to eNOS uncoupling^81^. It is conceivable cysteine oxidation or nitrosation by peroxynitrite in HFD could prevent palmitoylation. Because the main NO donor used for our experiments (DETA/NO) provides NO rather than peroxynitrite, our results provide strong evidence for NO delivering the nitrosation modification. In addition, most of the data presented here is in non-obese mice where eNOS uncoupling is not prevalent. It is also possible other parts of the pathway may also be nitrosated. For example, the biphasic effects of cysteine oxidation and nitrosation has recently been shown to regulate key endothelial proteins like vascular endothelial growth factor (VEGF)^82^ and Cav1^21, 23^. It is even possible eNOS itself may be both nitrosated and palmitoylated which could help move CD36 to and from the plasma membrane. Further work is warranted in these areas.

Last, elevated serum lipid levels are clinically characterized as hyperlipidemia which continues to be a crucial indicator of metabolic syndrome and cardiovascular disease risk. For patients with elevated cholesterol, statins are a first line prescription to manage the progression of hyperlipidemia by targeting the LDL receptor and the synthesis of cholesterol in the liver. However, up to 30% of patients on statins are intolerant and never reach healthy cholesterol levels^83^. This represents nearly 200 million people worldwide that have uncontrolled serum lipid levels. In addition, statin use does not come without risks, including increased risk of diabetes and damaged muscle. Thus, there is a need for a new lipid lowering modalities. CD36 is well-documented to regulate lipid uptake into cells but, crucial parts of this fatty acid translocases’ biology are not fully understood. Exploiting the mechanisms by which CD36 imports lipids could be a novel way to manage hyperlipidemia. Our work adds important mechanistic insight into how the properties of CD36 could be targeted for drug development by focusing on balancing the post-translational modification of palmitoylation and nitrosation

## Materials and Methods

### Animals

Only male mice were used, 20-23 weeks of age, on a C57Bl/6 genetic background, and were cared for under the provisions of the University of Virginia Animal Care and Use Committee and followed the National Institutes of Health guidelines for the care and use of laboratory animals. Animals were subject to a 12hr light dark cycle. Mice were housed on standard corn cobb bedding expect for metabolic studies where mice were fasted overnight on wood chip bedding. A C57BL/6n mice were purchased from Taconic. The inducible, EC-specific Cav1 knockout mice (Cdh5ER^T2+^/*Cav1^fl/fl^*) were generated by crossing Cdh5ER^T2+^/*Cav1^WT/WT^* mice (a kind gift from Dr Ralf Adams, Max Plank Institute, Germany) with Cdh5ER^T2-^/*Cav1^fl/fl^* mice^79, 84, 85^. To conditionally induce Cav1 deletion in the vascular endothelium, Cdh5ER^T2-^/*Cav1^fl/fl^* (EC Cre^-^ *Cav1^fl/f^*) and Cdh5ER^T2+^/*Cav1^fl/fl^* (EC Cre^+^ *Cav1^fl/fl^*) littermates received intraperitoneal (I.P.) injections of Tamoxifen (1 mg in 0.1 ml peanut oil) at six weeks of age for 10 consecutive days. For diet studies, littermates were randomly assigned to normal chow (NC; 5% Kcal from fat, house chow) or high fat diet (HFD; 60% Kcal from fat Bio-Serv-The Foster Corp F3282) starting at 8 weeks of age for 12 consecutive weeks. Additionally, C57BL6J (strain #: 000664) and eNOS transgenic mice (strain#: 002684) were purchased from Jackson Laboratories and used between the ages of 20-22 weeks. The *eNOS^fl/fl^* mice on both the endothelial and red blood cell (RBC) cre were used as described in^22^. Age of all mice at the completion of NC or HFD are 20 weeks. For L-NAME administration to NC mice, L-NAME (98%, Fisher Scientific: AAH6366606) was provided in the drinking water at 60mg/dL for 14 days. For L-NAME administration to HFD mice, L-NAME was provided in the drinking water at 100mg/dL for 14 days. All experiments were performed on a minimum of four mice. For all assessments of blood, blood was collected via terminal cardiac puncture using a syringe fitted with 25G needle, coated with EGTA to prevent clotting and deposited in gold cap blood collection tubes. Blood lipids (Cholesterol, LDL, HDL, and triglycerides) were processed by UVA clinical laboratory. Serum nitrite and nitrate levels were measured as described in Leo et al. 2021^22^. Non-esterified fatty acids (NEFA) were measured using Fujifilm NEFA absorbance kit (Fujifilm Medical systems: 633-52001). Diet and experimental procedures for ROSA26-eYFP^+/+^ Cdh5-CreER^T2+^ mice used for scRNAseq are described in depth in Dunaway and Luse et al. 2023.^40^

### Genomic excision

DNA was extracted from lung tissue and digested using proteinase K (Bioline: BIO-37084) for genomic excision gels. A set of three primers were used F: TTCTGTGTGCAAGCCTTTCC, R1: GTGTGCGCGTCATACACTTG, R2: GGGGAGGAGTAGAAGGTGGC. Non-excised Cav1 mice have a band at ∼7Kb and Cav1 excision is seen as an absence of a band.

### Tissue culture

Human adipose microvascular endothelial cells (HAMECs) were acquired from ScienCell (#7200) and cultured according to supplier guidelines. All HAMECs used were under passage 10. mRNA knockdown was performed using siRNA; Cav1 (30nM, Origene: SR319567) and CD36 (10nM, Thermo Fisher: s534752). 10mM Dimethyl-b-cyclodextrin (Thermo: 235530050) was used to deplete HAMECs of cholesterol. Knockdown of Cav1 was achieved via nucleofection using a LONZA nucleofector^®^, while lipofectamine (Thermo Fisher: L3000008) was used for knockdown of CD36. Briefly, cells were washed with PBS and incubated in serum free media containing 10mM dimethyl-b-cyclodextrin at 37°C for 1hr. Cholesterol depletion was validated using a Cholesterol/Cholesterol ester-glo assay (Promega: J3190). Endocytosis was blocked using two drugs Dynosore (selleckchem: S8047) and Genistein (Fisher: G02721) at 80µM and 200µM respectively for 30min at 37°C. For lipid treatment experiments, cells were treated with 50µM lipids (12:5µM linoleic acid, 25µM oleic acid, 12.5µM palmitic acid). Briefly, lipid stocks were made in ethanol and for each experiment the appropriate volume of lipids were dried down using nitrogen gas and then resuspended in cell culture media. The media was then sonicated and vortexed three times each and allowed to warm to 37°C. Lipid media was passed through a 0.45µM filter before being added to cells for 30 minutes. All cells to be imaged were washed once with 1X PBS and then fixed with 4% paraformaldehyde for 10 minutes at room temperature. Cellular lipid staining was accomplished through the use of BODIPY493/503 (Thermo Fisher: D3922) dissolved first in DMSO at a concentration of 1mg/mL then further diluted in PBS to fixed cells at 1µM. Cells were treated with L-NAME at concentration of 1mMol for 18hours either 72hrs after a control or siCav1 transfection. HAMECs were transfected with mCherry-CD36-C-10 (Addgene: Plasmid #55011), 8µg of plasmid was used per 1 million cells using nucleofection.

HEK293T were cultured in DMEM, high glucose (Gibco, 11965-092) supplemented with 1% Pen-Strep (Gibco, 15140122), 10% FBS (Avantor: 97068-085). Cells were transfected with mCherry-CD36-C-10 (Addgene: Plasmid #55011), 5µg per 10cm dish or 0.5µg per well of a 6-well dish, using Lipofectamine 3000 (Thermo Fisher: L3000001). Point mutations to the mCherry-CD36-C-10 plasmids were generated with the help of Genscript. HEK293T cells were treated with NO donors either DETA/NO-NONoate (DETA/NO: 50µM 30 minutes, Fisher: AC328650250) or S-Nitrosoglutathione (SNOG: 100µM 30 minutes, Fisher: 06-031-0) for nitrosation experiments. HEK293T cells treated with lipids followed the same procedure state above for HAMECs. Cells were fixed for 10min at room temperature with 4% paraformaldehyde and then stained with bodipy in PBS at 1µM.

### Lipid raft fractionation

HAMECs were homogenized in 25mM MES and 150mM NaCl then centrifuged at 1000g for 10min at 4°C. Supernatant was collected and mixed with various percentages of Optiprep (Sigma: D1556) to create a discontinuous density gradient. Samples were centrifuged at 380 x 1000 RPM for 2 hours at 4°C in a Beckman ultracentrifuge. Post centrifugation fractions were removed from top to bottom to isolate lipid rafts based on density. Lipid raft fractionation was performed 72hrs after SiCav1 or control treatment.

### Biotin Switch

Biotin switch assay was performed using the S-Nitrosylated Protein Detection Kit (Cayman chemical: 10006518) and HEK293T cells. Briefly, cells were washed with supplied wash buffer and free thiols were blocked. Samples were split into two providing internal controls where one sample did not receive the reducing and detection agents (biotin). After biotin was added to thiols containing nitrations, streptavidin beads (Thermo Fisher: 65601) were used to isolate biotinylated/nitrated proteins. Protein bound to the beads was eluted using 5x laemmli buffer at room temperature for 15 minutes and then heated at 95°C for 1 minutes. Sample was loaded according to western blotting procedures (below) and immunoblotted for CD36 (Thermo fisher PA1-16913) or mcherry (abcam: ab213511) tag on CD36.

### Real-time quantitative PCR

Total RNA was extracted from mouse tissues using the Aurum Total RNA Fatty and Fibrous Tissue Extraction kit (Biorad: #732-6870). RNA from cells was extracted using Zymo Research R1055 Quick-RNA MiniPrep Kit, Zymo Research Kit (Genesee: 11-328). RNA concentration was measured using the Nanodrop1000 spectrophotometer (Thermo Fisher). RNA was stored at -80°C before reverse transcription with SuperScript III First-Strand Synthesis system (Thermo Fisher: 18080051) using random hexamer primers on 1μg of template RNA. Real-time quantitative PCR was performed using Taqman Gene Expression Master Mix (Thermo Fisher: 4369016) and Taqman Real-Time PCR assays in MGB-FAM for Cav1 (Hs00971716_m1; Mm00432403_m1), CD36 (Hs00354519_m1; Mm00432403_m1), and were normalized to β-2-microglobin/B2M in VIC-PL (Hs00364808_m1; Mm00437762_m1). Reactions were run in a CFX Real-Time Detection System (Applied BioSystems) and threshold cycle number (CT) was used as part of the 2-DDCT method to calculate fold change from control.

### Western Blotting

Cell and tissue lysates were generated in RIPA (50mmol/L Tris-HCL, 150mmol/L NaCl, 5mmol/L EDTA,1% deoxycholate, 1% Triton-X100) in PBS and pH adjusted to 7.4) supplemented with protease inhibitor cocktail (Sigma: P8340). Lysates were rocked at 4°C for 30-60 min to solubilize proteins, sonicated briefly, and centrifuged for 15 min at 12,000 rpm to pellet cell debris. Protein concentration was determined using the Pierce BCA method (Thermo Fisher: 23227). 20μg of total protein was loaded into each sample well. Samples were subjected to SDS gel electrophoresis using 8% or 4-12% Bis-Tris gels (Invitrogen) and transferred nitrocellulose membranes for immunoblotting. Membranes were blocked for 1 hour at room temperature in a solution containing 3% BSA in Tris buffered saline, then incubated overnight at 4°C with primary antibodies against Cav1 (abcam: ab32577, 1:2000), CD36 (Human tissue Thermofisher: PA1-16813 1:500, mouse tissue abcam: ab124515 1:500), Calnexin (abcam: ab22595 1:1000), and eNOS (BD transduction: 610297 1:1000). Membranes were washed and incubated in LiCOR IR Dye secondary antibodies (1:10,000) for 1 hour and viewed/quantified using the LiCOR Odyssey CLx with Image Studio software. Licor Total Protein stain was used for loading normalization. Representative western blot images have been cropped for presentation.

### Single-Cell RNA sequencing (scRNAseq)

A previously published scRNA-seq dataset from human adipose tissue was used to create these data^41^. Endothelial cells were subsetted, using subset() function, from previously published data using clusters defined by the original authors and confirmed by our arterial, venous, capillary, and lymphatic endothelial markers. Endothelial cells from visceral adipose tissue were used for analysis for more appropriate comparison with epididymal adipose ECs.

scRNAseq from mouse adipose endothelium has been previously described in^40^. Briefly, the generation of single cell indexed libraries was performed by the School of Medicine Genome Analysis and Technology Core, RRID:SCR_018883, using the 10X Genomics chromium controller platform and the Chromium Single Cell 3′ Library & Gel Bead Kit v3.1 reagent. Around 5,000 cells were targeted per sample and loaded onto each well of a Chromium Single Cell G Chip to generate single cell emulsions primed for reverse transcription. After breaking the emulsion, the single cell specific barcoded DNAs were subjected to cDNA amplification and QC on the Agilent 4200 TapeStation Instrument, using the Agilent D5000 kit. A QC run was performed on the Illumina Miseq using the nano 300Cycle kit (1.4 Million reads/run), to estimate the number of targeted cells per sample using the Cellranger 3.0.2 function. After run completion, the Binary base call (bcl) files were converted to fastq format using the Illumina bcl2fastq2 software raw reads in fastq files were mapped to the mm10 reference murine genome and assigned to individual cells by CellRanger 5.0. Data from two separate experiments representing cells from 12 mice were analyzed in RStudio (2022.07.1) with the Seurat package (4.3.0). Sequencing yielded 27,944 cells with 54,100 features. To ensure high quality data, cells were excluded if they contained less than 200 genes, more than 5000 genes, if their transcriptome was more than 5% mitochondrial encoded, and more than 5% hemoglobin beta. This resulted in a final data set of 17,164 cells. Data was combined using SCTransform, normalized, and 3000 variable features were chosen. UMAPs were generated using 20 principal components. Clusters were generated using a resolution of 1. Non-endothelial cells were excluded based on low expression of Pecam1 and Cdh5 as well as high expression of non-endothelial markers (Col1a1, Acta2, Cd3g, Ptprc, Ccr5, Adipoq).

### En Face Imaging

Epididymal fat pad arteries (100-200µm in diameter) were collected and stripped of adipose and connective tissue. Arteries were subsequently fixed in 4% paraformaldehyde on ice for 10 minutes. For *en face* preparation, arteries were cut longitudinally with microdissection scissors and pinned open on polymerized Sylgard 184 (Electron Microscopy Sciences) using tungsten wire (0.0005”, ElectronTubeStore). For immunostaining, vessels were then permeabilized with 0.5% TritonX100 in PBS for 30 minutes at room temperature and blocked in 1% bovine serum albumin, Fraction V (BSA Sigma) in 0.5% TritonX100/PBS. Primary antibody staining was performed overnight at 4°C in 0.1% BSA in 0.5% TritonX100/PBS. Cav1 (Abcam: ab211503, 1:200), CD36 (abcam: ab124515, 1:100), Calnexin (abcam: ab219644, 1:100) primary antibodies were used. Samples were incubated with secondary antibodies at 1:500 for 1-2 hours at room temperature. For Nile red staining, samples were immediately stained after pinning. Nile red (Thermo: N1142) was used at 5µM in PBS for 30min at room temperature. Nuclei were stained using DAPI (Invitrogen: D1306, final concentration 0.1mg/mL) in addition to mounting with Prolong Gold Antifade Mountant (Invitrogen: P36930). Images were collected using an Olympus FV3000 with a 60X oil emersion lens and post processing was completed using FIJI.

### Mitochondrial Stress Test

HAMECs were plated (80,000 cells per well) in complete medium in XFe96 cell culture microplate and allowed to settle for one hour at room temperature before overnight culture. Sixteen hours prior to assessment HAMECs were treated with L-NAME (see above) and subsequently four hours prior to assessment HAMECs were treated with 50 μM lipids as well as DETA/NO (see above). Immediately prior to the assay the media was changed to mitochondrial stress test medium (Corning: 50-003-PB) including L-NAME or DETA/NO as indicated. Mitochondrial activity was assessed by measurement of O_2_ consumption rate on a Seahorse XFe96 instrument (Agilent Technologies). The rate of O_2_ change was measured every 13 minutes for a 3-minute interval before sequential challenge with 1) 1 μM oligomycin (Sigma-Aldrich: 75351), 2) 2 μM BAM15^86^ (Cayman Chemical Company: 17811) and 3) 10 μM antimycin A (Sigma-Aldrich: A8674) and 10 μM rotenone (Sigma-Aldrich: R88751G). Maximal OCR was measured as post-BAM15 OCR minus post-antimycin A/rotenone OCR. Data represented here have been normalized (pmol O_2_/μg total protein) to total protein in each well of assay plate using BCA assay (Thermo: 23227).

### RNA sequencing

ECs were treated as described above with the same methods used for “Mitochondrial stress test”. RNA was isolated using Quick-RNA MiniPrep Kit (Genesee Scientific: 11-328). When necessary, samples were further concentrated and purified with the RNeasy MinElute Cleanup Kit (Qiagen: 74804). PE150 reads were generated with the Illumina NovoSeq platform. The paired-end reads were mapped to the hg38 reference genome using STAR (v 2.7.9a). Gene counts were generated with FeatureCounts in the Rsubread (v. 2.8.8) package. DESeq2 (v. 1.34.0) was used to calculate log fold change and adjusted p-values. The adjusted p-values were then adjusted for multiple comparisons between groups by the Benjamini-Hochberg procedure using p.adjust(). Ensemble IDs were converted to gene symbols using the EnsDb.Hsapiens.v75 and org.Hs.eg.db packages. For gene set enrichment analysis, genes were ranked by log2foldchange and pathway analysis was conducted with clusterprofiler (v 4.2.2). Raw data and normalized counts have been deposited in NCBI’s Gene Expression Omnibus and are accessible through the GEO accession (GSE260897).

### Pressure Myography

Third order mesenteric arteries (diameter between 100-200µm) were removed from 14-week old C57BL/6J mice. Arteries were cannulated on two glass pipettes tips and equilibrated to 80mmHg as previously described^87^. For vasoreactivity measurements vessels were either incubated with 50µM L-NAME or water for 20 minutes. Subsequently for vessels receiving lipid treatments 10µM lipids (10µM oleic acid solubilized in ethanol) were added to the circulating buffer for an additional 20 minutes. Vessels were pre-constricted with 1µM phenylephrine (PE) and then exposed to serial doses of Acetylcholine (ACh) to assess vasodilation. Three male mice were used for the experiments using lipids with 1-2 vessels per mouse averaged for each sample value. Five mice were used for experiments without lipids with 1-2 vessels averaged per each sample value.

### Acyl–biotinyl exchange and Western blotting to analyze palmitoylation

Palmitoylation analysis was done using acyl–biotinyl exchange (ABE) as previously described^88^. One 10 cm plate with HEK-293 cells at 80-90% confluence was lysed in 2% SDS-containing buffer and free cysteines were blocked by 10 mM NEM. Then, palmitoylated cysteines were liberated by 0.4 M hydroxylamine and labeled with biotin-HPDP (Pierce). Biotinylated proteins were pulled down using streptavidin-magnetic beads (Dynabeads, Thermo Scientific) and eluted with 1% 2-mercaptoethanol. Three chloroform/methanol precipitations were performed between each step to remove chemicals. After elution, a Western blot of the eluate (palmitoylated fraction) and input (total protein) was performed for the different Rab proteins (endogenous) or TMD probes (anti-RFP). For quantification, densitometry analysis was performed in BioRAD ChemiDoc MP.

### Statistical Analysis

Statistical analysis was performed using Prism GraphPad 9 and R studio for scRNAseq and bulk RNA sequencing analysis. Specific statistical test type and N values are listed in each corresponding figure legend. Data represented are the mean with SEM (+ and -) shown as error bars. All N values represent different experiments (*in vitro*) and individual mice (*in vivo*). For image quantification, three fields of view were used to average each N value.

## Acknowledgments

We thank Anne Kenworthy (University of Virginia) for contributing her expertise on caveolin-1 and its role in caveolae and lipid nanodomain organization. We thank the University of Virginia School of Medicine Research Histology Core and Blood lab.

## Author contributions

Conceptualization: BEI, MAL, MK, NL

Methodology: PB, AK, TS, IL, RM

Investigation: MAL, WJS, LSD, SN, SKL, AC, RT, CP, TSK, XS, CAR, LC, KRL, MK, RM

Visualization: LC

Supervision: BEI

Writing—original draft: MAL, BEI

Writing—review & editing:

## Data and materials availability

All data are available in the main text or the supplementary materials

## Supplementary Materials

**Supplementary Figure 1:**
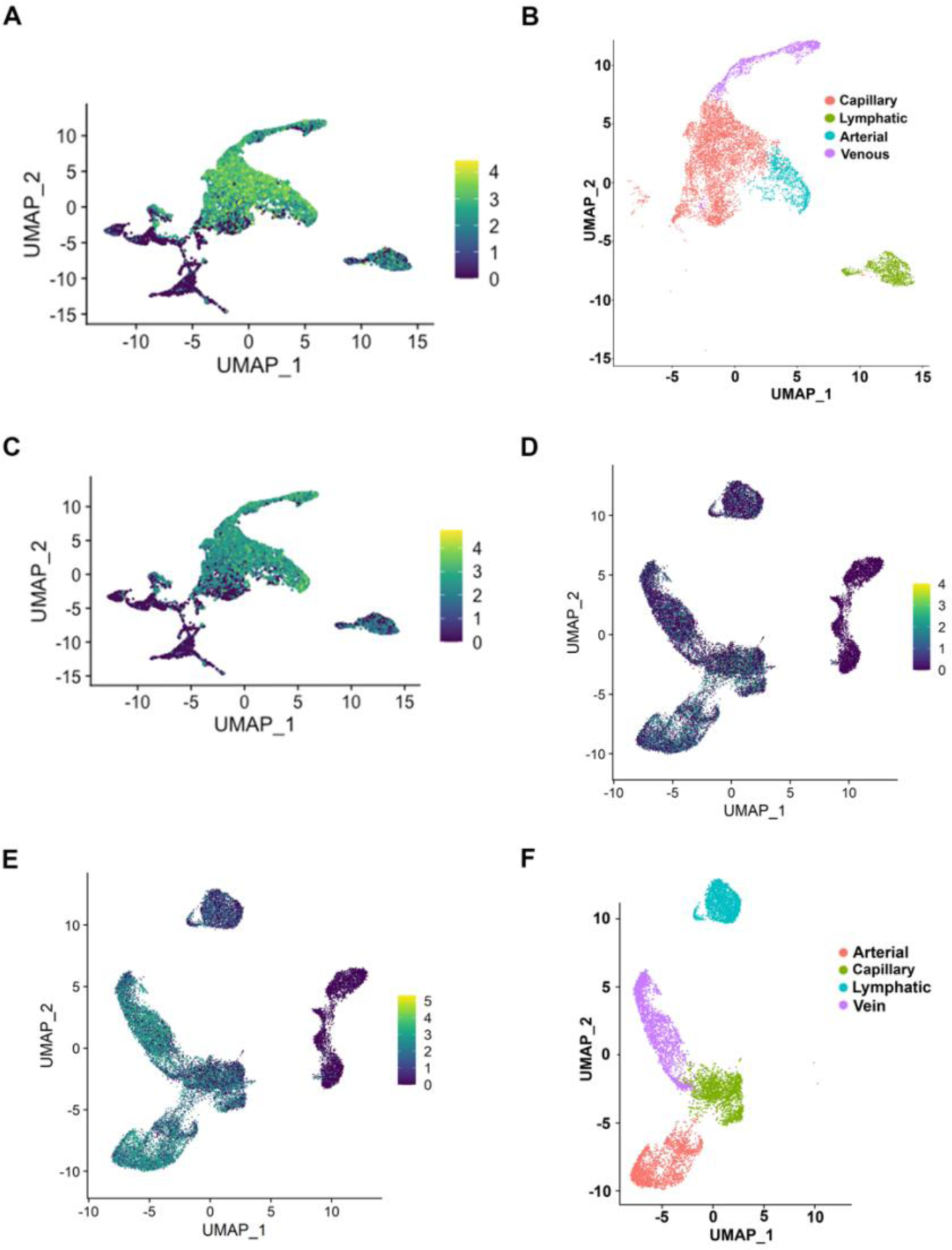
Filtering and subsetting scRNAseq clusters for EC enrichment. UMAP of all cells (lean and obese patients) in Emont data set showing expression pattern of CDH5 (**A**) and PECAM1 (**B**). (**C**) Filtered human adipose ECs clustered by vascular origin. UMAP of all adipose ECs (from HFD and NC mice) before filtration with expression patterns of Cdh5 (**D**) and Pecam1 (**E**). (**F**) Filtered murine adipose ECs clustered by vascular origin.

**Supplementary Figure 2:**
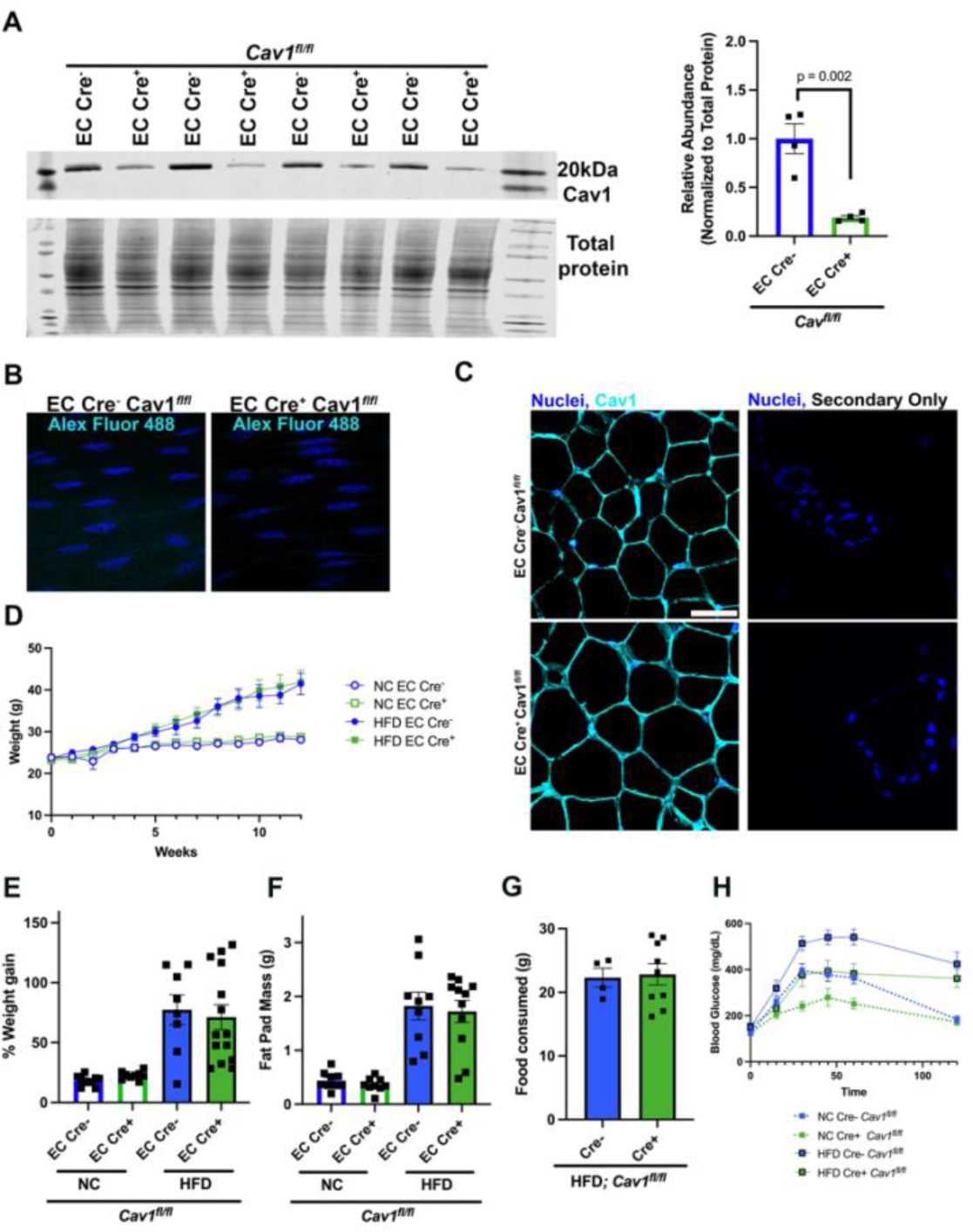
Caveolin 1 deletion from endothelial cells does not alter body mass. (**A**) Western blot from lung, showing loss of Cav1 protein when EC Cre is present, quantification on left is normalized to total protein loading control. Unpaired t-test was used, N = 4-5. (**B**) Adipose artery en face with secondary only staining. (**C**) Adipose tissue cross section stained with Cav1, Scale bar = 50µm. (**D**) Weight of mice (in grams; g) over the course of the 12-week NC or HFD diets of *Cav1^fl/fl^* mice (**E**) Percent weight gain calculated using ((final mass-initial mass/Initial mass)*100 of high fat diet (HFD) and normal chow (NC) mice. (**F**) Epididymal fat mass and food intake (**G**) of EC Cre^-^ (N.=4) and EC Cre+ (N= 9) *Cav1^fl/fl^* mice. (**H**) Glucose tolerance test, blood glucose over time after glucose injection. N= 8-10 mice for all experiments for E,F, and H.

**Supplementary Figure 3:**
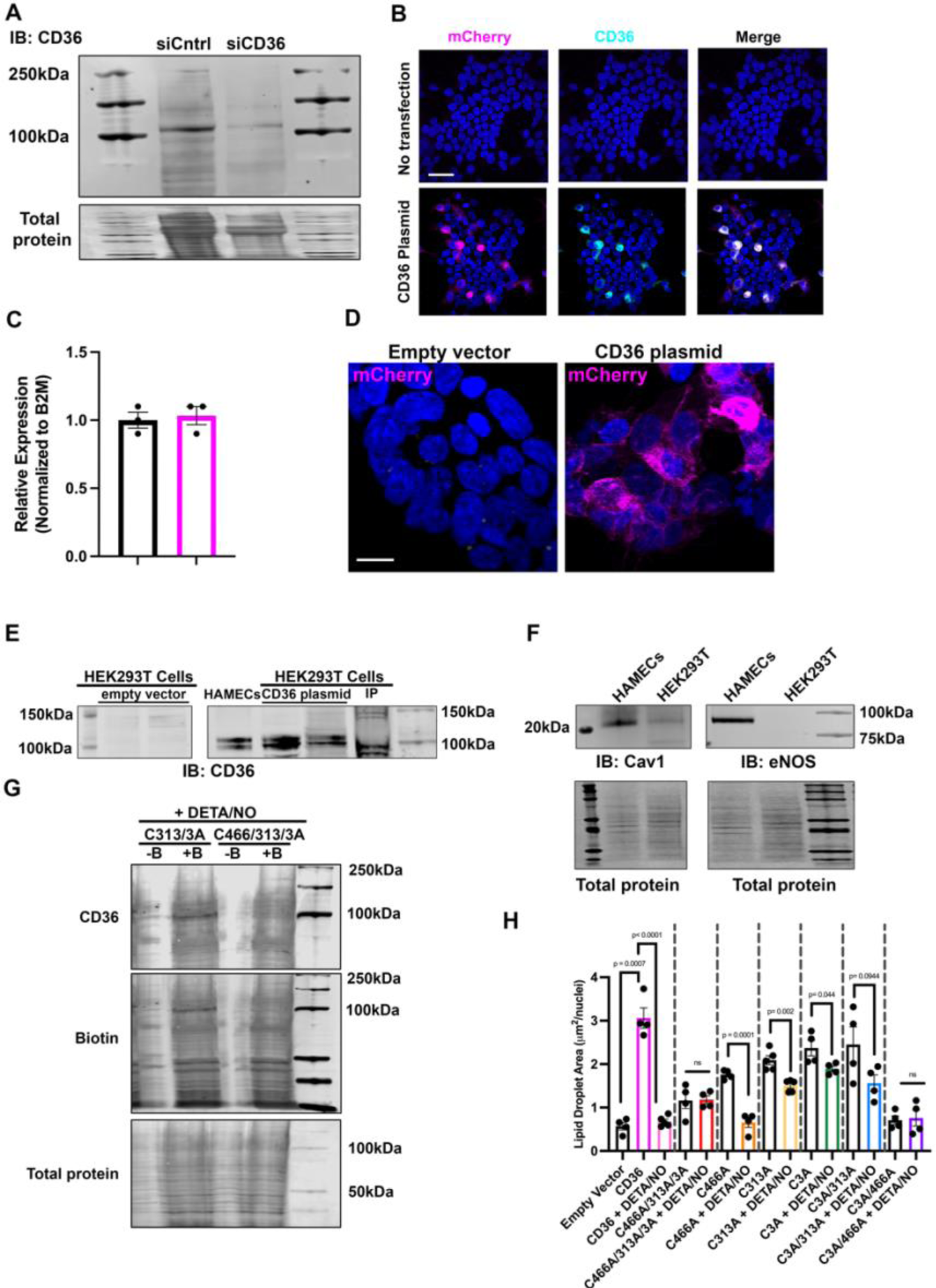
Protein composition of HAMEC and HEK cells. (A) Western blot showing loss of CD36 band with siCD36 in HAMECs, below is total protein **(B)** HEK cells with no transfection (top) and HEK cells transfected with CD36 plasmid (bottom). CD36 plasmid is tagged with mCherry (**C**) Cav1 RNA expression with CD36 knockdown in HAMECs measure via q-PCR. (**D**) HEK cells transfected with CD36 plasmid with mcherry reported. (**E**) HEK cell transfections with CD36 plasmid with subsequent CD36 immunoblot. IP = immunoprecipitation. (**F**) Cav1 and eNOS immunoblots in HAMEC vs. HEK cells. (**G**) Biotin switch assay to detect nitrosation of CD36 using C313/3A and C466/313/3A mutant versions of the CD36 plasmid. -B an +B represent without or with biotin respectively. All samples were treated with DETA/NO. (**H**) Cumulative HEK293T lipid uptake data.

**Supplementary Figure 4:**
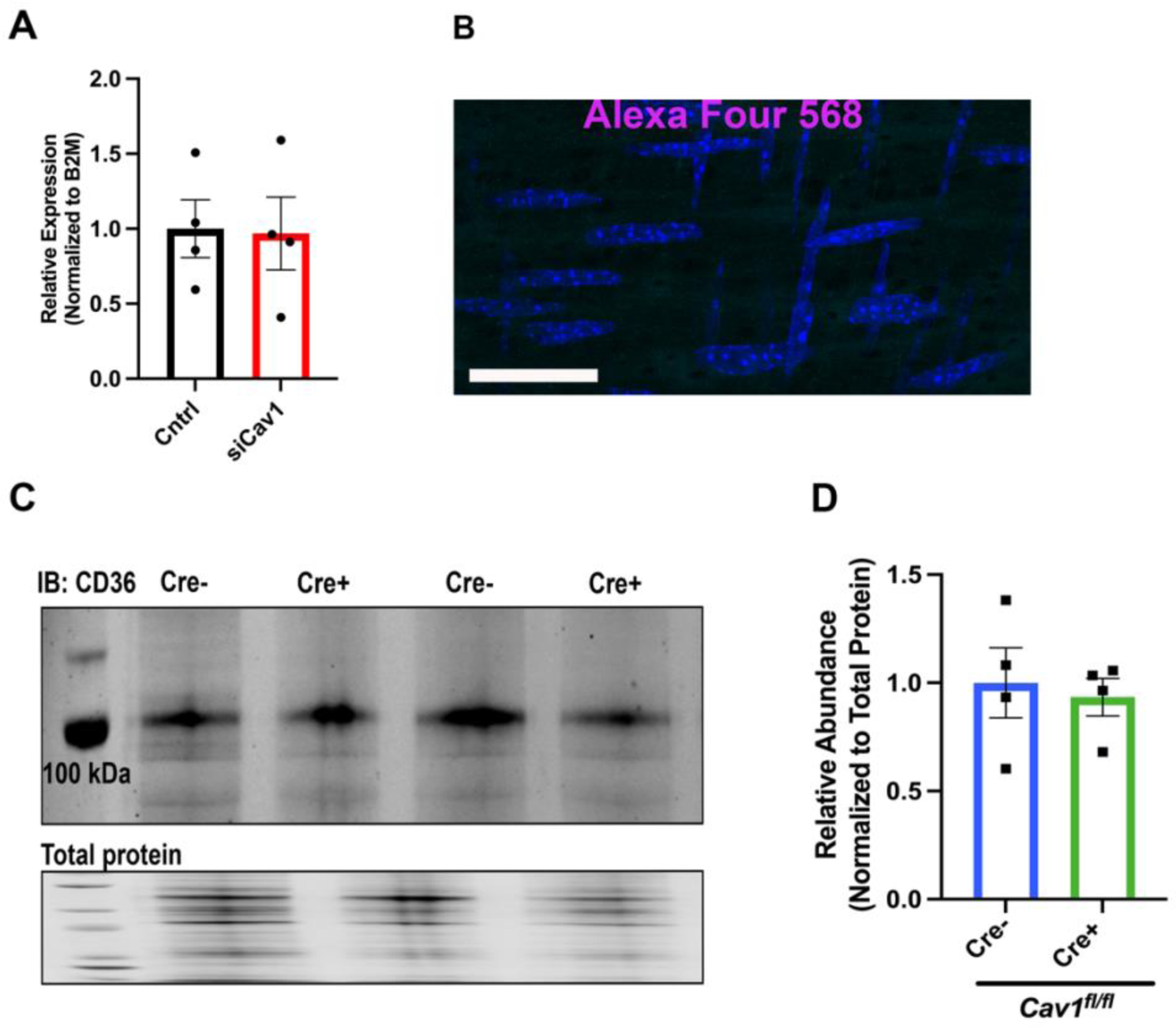
CD36 protein and RNA levels are unaltered with loss of Caveolin 1. (**A**) CD36 expression in control and siCav1 treated cells measured via q-PCR. (**B**) Adipose artery en face stained with Alexa Four 568 only. (**C**) CD36 immunoblot from EC Cre^-^ and EC Cre^+^ *Cav1^fl/fl^* mice and quantified in (**D**) by normalizing to total protein. N = 4 mice.

**Supplementary Figure 5:**
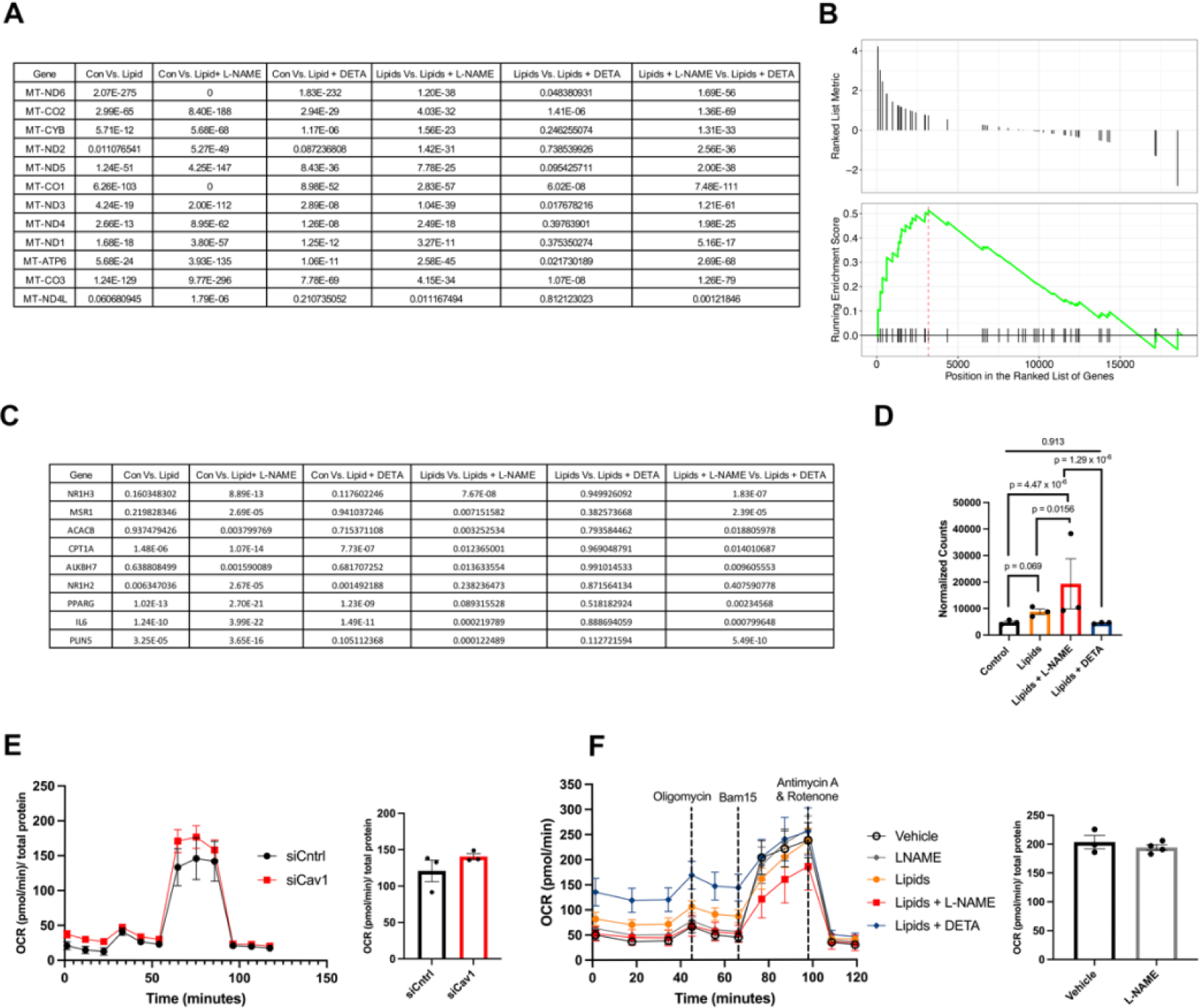
Oxygen consumption and mitochondrial gene expression is decreased with lipids. (**A**) P values of mitochondrial genes, using DESeq2 and the Benjamini-Hochberg correction as described in the methods. (**B** and **C**) Gene set enrichment analysis (GSEA) for regulation of lipid storage genes. GSEA plot for Lipid + L-NAME treated HAMECs. P values for lipid storage genes. (**D**) HAMEC *FABP4* expression were calculated using DESeq2 and the Benjamini-Hochberg correction as described in the methods. N= 3 for all bulk RNA sequencing data. (**E**) Oxygen consumption trace for mitochondrial stress test measured via seahorse assay. siCntrl (black) or siCav1 (red) HAMECs were used. Maximum oxygen consumption (right) and Oxygen consumption over time normalized to total cellular protein. (**F**) Mitochondrial stress test for HAMECs treated with L-NAME (grey), Lipids (orange), Lipids + L-NAME (red), Lipids + DETA/NO (blue) or control (black) media. Maximum oxygen consumption for control and L-NAME treated cells (right).

**Supplementary Figure 6:**
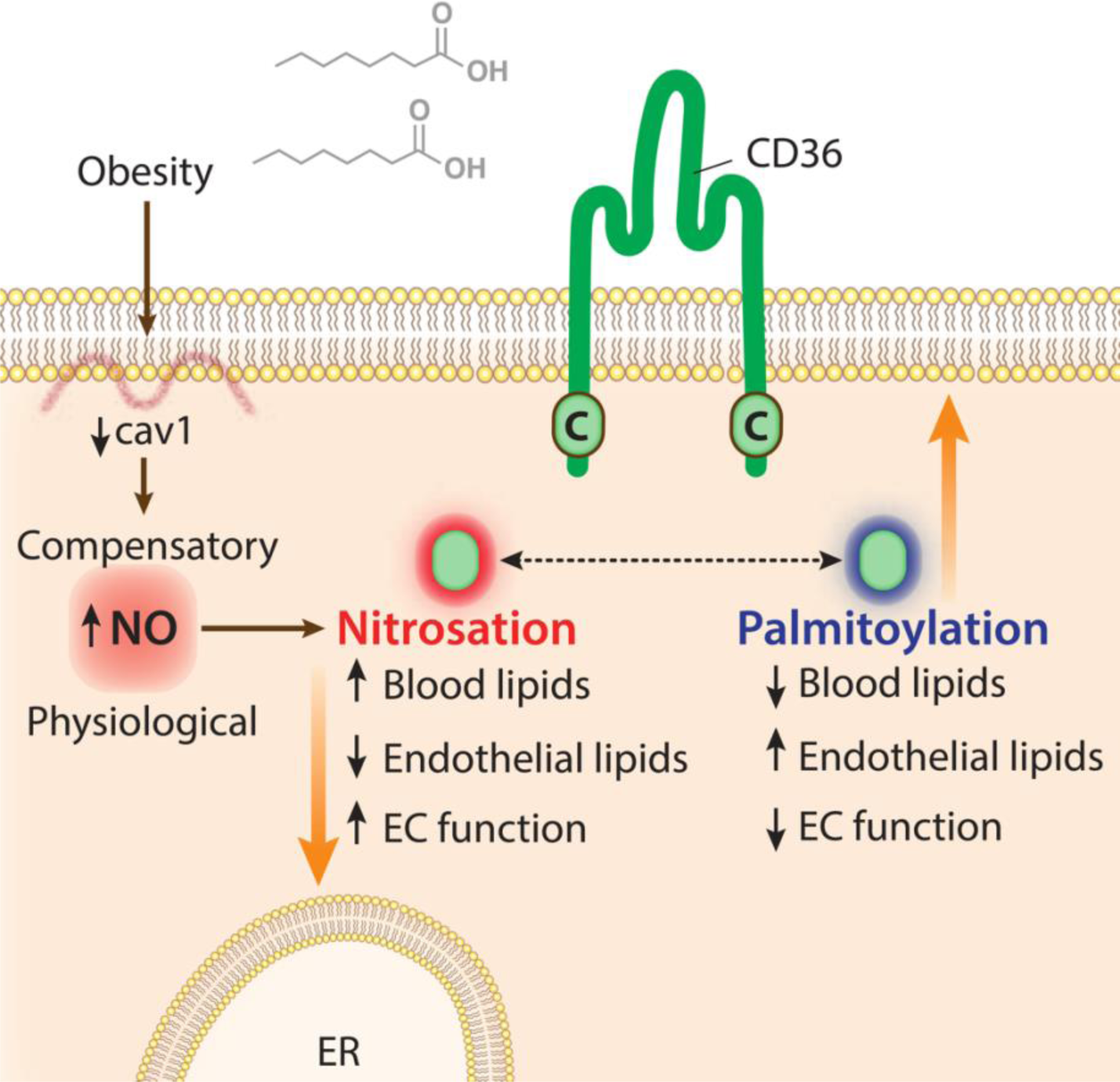
Summary figure showing proposed mechanism for nitrosation-palmitoylation of CD36 which regulates endothelial lipid uptake.

## References

1. Mehrotra D, Wu J, Papangeli I and Chun HJ. Endothelium as a gatekeeper of fatty acid transport. Trends in Endocrinology & Metabolism. 2014;25:99–106.

2. Yazdani S, Jaldin-Fincati JR, Pereira RV and Klip A. Endothelial cell barriers: Transport of molecules between blood and tissues. Traffic. 2019;20:390–403.

3. Wang H, Liu Z, Li G and Barrett EJ. The vascular endothelial cell mediates insulin transport into skeletal muscle. American Journal of Physiology-Endocrinology and Metabolism. 2006;291:E323–E332.

4. Kuo A, Lee MY and Sessa WC. Lipid droplet biogenesis and function in the endothelium. Circulation research. 2017;120:1289–1297.

5. Stoletov K, Fang L, Choi S-H, Hartvigsen K, Hansen LF, Hall C, Pattison J, Juliano J, Miller ER and Almazan F. Vascular lipid accumulation, lipoprotein oxidation, and macrophage lipid uptake in hypercholesterolemic zebrafish. Circulation research. 2009;104:952–960.

6. Jang C, Oh SF, Wada S, Rowe GC, Liu L, Chan MC, Rhee J, Hoshino A, Kim B and Ibrahim A. A branched-chain amino acid metabolite drives vascular fatty acid transport and causes insulin resistance. Nature medicine. 2016;22:421–426.

7. Listenberger LL, Han X, Lewis SE, Cases S, Farese Jr RV, Ory DS and Schaffer JE. Triglyceride accumulation protects against fatty acid-induced lipotoxicity. Proceedings of the National Academy of Sciences. 2003;100:3077–3082.

8. Unger RH and Scherer PE. Gluttony, sloth and the metabolic syndrome: a roadmap to lipotoxicity. Trends in Endocrinology & Metabolism. 2010;21:345–352.

9. Boutagy NE, Gamez-Mendez A, Fowler JW, Zhang H, Chaube BK, Esplugues E, Lee S, Horikami D, Zhang J and Citrin KM. Dynamic metabolism of endothelial triglycerides protects against atherosclerosis in mice. The Journal of Clinical Investigation. 2024.

10. Boutagy NE, Singh AK and Sessa WC. Targeting the vasculature in cardiometabolic disease. The Journal of Clinical Investigation. 2022;132.

11. Simionescu M. Implications of early structural-functional changes in the endothelium for vascular disease. Arteriosclerosis, thrombosis, and vascular biology. 2007;27:266–274.

12. Fernández-Hernando C, Yu J, Suárez Y, Rahner C, Dávalos A, Lasunción MA and Sessa WC. Genetic evidence supporting a critical role of endothelial caveolin-1 during the progression of atherosclerosis. Cell metabolism. 2009;10:48–54.

13. Fernández-Hernando C, Yu J, Dávalos A, Prendergast J and Sessa WC. Endothelial-specific overexpression of caveolin-1 accelerates atherosclerosis in apolipoprotein E-deficient mice. The American journal of pathology. 2010;177:998–1003.

14. Wang H, Wang AX and Barrett EJ. Caveolin-1 is required for vascular endothelial insulin uptake. American Journal of Physiology-Endocrinology and Metabolism. 2011;300:E134–E144.

15. Kakava S, von Eckardstein A and Robert J. Regulation of low-density lipoprotein transport through endothelial cells by caveolae. Atherosclerosis. 2023;375:84–86.

16. Le Master E, Paul A, Lazarko D, Aguilar V, Ahn SJ, Lee JC, Minshall RD and Levitan I. Caveolin-1 is a primary determinant of endothelial stiffening associated with dyslipidemia, disturbed flow, and ageing. Scientific Reports. 2022;12:17822.

17. Shamsaldeen YA, Ugur R, Benham CD and Lione LA. Diabetic dyslipidaemia is associated with alterations in eNOS, caveolin-1, and endothelial dysfunction in streptozotocin treated rats. Diabetes/Metabolism Research and Reviews. 2018;34:e2995.

18. Li X, Sun X and Carmeliet P. Hallmarks of endothelial cell metabolism in health and disease. Cell metabolism. 2019;30:414–433.

19. Matthaeus C, Lahmann I, Kunz S, Jonas W, Melo AA, Lehmann M, Larsson E, Lundmark R, Kern M and Blüher M. EHD2-mediated restriction of caveolar dynamics regulates cellular fatty acid uptake. Proceedings of the National Academy of Sciences. 2020;117:7471–7481.

20. Bernatchez P, Sharma A, Bauer PM, Marin E and Sessa WC. A noninhibitory mutant of the caveolin-1 scaffolding domain enhances eNOS-derived NO synthesis and vasodilation in mice. The Journal of clinical investigation. 2011;121.

21. Chen Z, DS Oliveira S, Zimnicka AM, Jiang Y, Sharma T, Chen S, Lazarov O, Bonini MG, Haus JM and Minshall RD. Reciprocal regulation of eNOS and caveolin-1 functions in endothelial cells. Molecular Biology of the Cell. 2018;29:1190–1202.

22. Leo F, Suvorava T, Heuser SK, Li J, LoBue A, Barbarino F, Piragine E, Schneckmann R, Hutzler B and Good ME. Red blood cell and endothelial eNOS independently regulate circulating nitric oxide metabolites and blood pressure. Circulation. 2021;144:870–889.

23. Bakhshi FR, Mao M, Shajahan AN, Piegeler T, Chen Z, Chernaya O, Sharma T, Elliott WM, Szulcek R and Bogaard HJ. Nitrosation-dependent caveolin 1 phosphorylation, ubiquitination, and degradation and its association with idiopathic pulmonary arterial hypertension. Pulmonary circulation. 2013;3:816–830.

24. Son N-H, Basu D, Samovski D, Pietka TA, Peche VS, Willecke F, Fang X, Yu S-Q, Scerbo D and Chang HR. Endothelial cell CD36 optimizes tissue fatty acid uptake. The Journal of clinical investigation. 2018;128:4329–4342.

25. Abumrad NA, Cabodevilla AG, Samovski D, Pietka T, Basu D and Goldberg IJ. Endothelial cell receptors in tissue lipid uptake and metabolism. Circulation research. 2021;128:433–450.

26. Peche V, Pietka T, Jacome-Sosa M, Samovski D, Palacios H, Chatterjee-Basu G, Dudley A, Beatty W, Meyer G and Goldberg I. Endothelial cell CD36 regulates membrane ceramide formation, exosome fatty acid transfer and circulating fatty acid levels. Nature communications. 2023;14:4029.

27. Dawson DW, Pearce SFA, Zhong R, Silverstein RL, Frazier WA and Bouck NP. CD36 mediates the in vitro inhibitory effects of thrombospondin-1 on endothelial cells. The Journal of cell biology. 1997;138:707–717.

28. Love-Gregory L, Sherva R, Sun L, Wasson J, Schappe T, Doria A, Rao DC, Hunt SC, Klein S and Neuman RJ. Variants in the CD36 gene associate with the metabolic syndrome and high-density lipoprotein cholesterol. Human molecular genetics. 2008;17:1695–1704.

29. Hao J-W, Wang J, Guo H, Zhao Y-Y, Sun H-H, Li Y-F, Lai X-Y, Zhao N, Wang X and Xie C. CD36 facilitates fatty acid uptake by dynamic palmitoylation-regulated endocytosis. Nature communications. 2020;11:4765.

30. Wang J, Hao J-W, Wang X, Guo H, Sun H-H, Lai X-Y, Liu L-Y, Zhu M, Wang H-Y and Li Y-F. DHHC4 and DHHC5 facilitate fatty acid uptake by palmitoylating and targeting CD36 to the plasma membrane. Cell reports. 2019;26:209–221. e5.

31. Tao N, Wagner SJ and Lublin DM. CD36 is palmitoylated on both N-and C-terminal cytoplasmic tails. Journal of Biological Chemistry. 1996;271:22315–22320.

32. Guan X and Fierke CA. Understanding protein palmitoylation: Biological significance and enzymology. Science China Chemistry. 2011;54:1888–1897.

33. Greaves J, Carmichael JA and Chamberlain LH. The palmitoyl transferase DHHC2 targets a dynamic membrane cycling pathway: regulation by a C-terminal domain. Molecular biology of the cell. 2011;22:1887–1895.

34. Zhang MM, Tsou LK, Charron G, Raghavan AS and Hang HC. Tandem fluorescence imaging of dynamic S-acylation and protein turnover. Proceedings of the National Academy of Sciences. 2010;107:8627–8632.

35. Salaun C, Greaves J and Chamberlain LH. The intracellular dynamic of protein palmitoylation. Journal of Cell Biology. 2010;191:1229–1238.

36. Kharitonov VG, Sundquist AR and Sharma VS. Kinetics of nitrosation of thiols by nitric oxide in the presence of oxygen. Journal of Biological Chemistry. 1995;270:28158–28164.

37. Stamler JS, Simon DI, Osborne JA, Mullins ME, Jaraki O, Michel T, Singel DJ and Loscalzo J. S-nitrosylation of proteins with nitric oxide: synthesis and characterization of biologically active compounds. Proceedings of the National Academy of Sciences. 1992;89:444–448.

38. Lohman AW, Weaver JL, Billaud M, Sandilos JK, Griffiths R, Straub AC, Penuela S, Leitinger N, Laird DW and Bayliss DA. S-nitrosylation inhibits pannexin 1 channel function. Journal of Biological Chemistry. 2012;287:39602–39612.

39. Gould N, Doulias P-T, Tenopoulou M, Raju K and Ischiropoulos H. Regulation of protein function and signaling by reversible cysteine S-nitrosylation. Journal of Biological Chemistry. 2013;288:26473–26479.

40. Dunaway LS, Luse MA, Nyshadham S, Bulut G, Alencar GF, Chavkin NW, Cortese-Krott MM, Hirschi KK and Isakson BE. Obesogenic diet disrupts tissue specific mitochondrial gene signatures in the artery and capillary endothelium. Physiological Genomics. 2023.

41. Emont MP, Jacobs C, Essene AL, Pant D, Tenen D, Colleluori G, Di Vincenzo A, Jørgensen AM, Dashti H and Stefek A. A single-cell atlas of human and mouse white adipose tissue. Nature. 2022;603:926–933.

42. Porta JC, Han B, Gulsevin A, Chung JM, Peskova Y, Connolly S, Mchaourab HS, Meiler J, Karakas E and Kenworthy AK. Molecular architecture of the human caveolin-1 complex. Science Advances. 2022;8:eabn7232.

43. Shu X, Keller TS, Begandt D, Butcher JT, Biwer L, Keller AS, Columbus L and Isakson BE. Endothelial nitric oxide synthase in the microcirculation. Cellular and molecular life sciences. 2015;72:4561–4575.

44. Ju H, Zou R, Venema VJ and Venema RC. Direct interaction of endothelial nitric-oxide synthase and caveolin-1 inhibits synthase activity. Journal of Biological Chemistry. 1997;272:18522–18525.

45. Murata T, Lin MI, Huang Y, Yu J, Bauer PM, Giordano FJ and Sessa WC. Reexpression of caveolin-1 in endothelium rescues the vascular, cardiac, and pulmonary defects in global caveolin-1 knockout mice. The Journal of experimental medicine. 2007;204:2373–2382.

46. Mahammad S and Parmryd I. Cholesterol depletion using methyl-β-cyclodextrin. Methods in membrane lipids. 2015:91–102.

47. Martens JR, Sakamoto N, Sullivan SA, Grobaski TD and Tamkun MM. Isoform-specific localization of voltage-gated K+ channels to distinct lipid raft populations: targeting of Kv1. 5 to caveolae. Journal of Biological Chemistry. 2001;276:8409–8414.

48. Seto S, Krishna S, Yu H, Liu D and Khosla S. Impaired Acetylcholine-Induced Endothelium-Dependent Aortic Relaxation by Caveolin-1 in. 2013.

49. Rees D, Palmer R, Schulz R, Hodson H and Moncada S. Characterization of three inhibitors of endothelial nitric oxide synthase in vitro and in vivo. British journal of pharmacology. 1990;101:746.

50. Silva FCd, Araújo BJd, Cordeiro CS, Arruda VM, Faria BQ, Guerra JFDC, Araújo TGD and Fürstenau CR. Endothelial dysfunction due to the inhibition of the synthesis of nitric oxide: Proposal and characterization of an in vitro cellular model. Frontiers in Physiology. 2022;13:978378.

51. Febbraio M and Silverstein RL. CD36: implications in cardiovascular disease. The international journal of biochemistry & cell biology. 2007;39:2012–2030.

52. Pohl J, Ring A, Korkmaz U, Ehehalt R and Stremmel W. FAT/CD36-mediated long-chain fatty acid uptake in adipocytes requires plasma membrane rafts. Molecular biology of the cell. 2005;16:24–31.

53. Ring A, Le Lay S, Pohl J, Verkade P and Stremmel W. Caveolin-1 is required for fatty acid translocase (FAT/CD36) localization and function at the plasma membrane of mouse embryonic fibroblasts. Biochimica et Biophysica Acta (BBA)-Molecular and Cell Biology of Lipids. 2006;1761:416–423.

54. Gerbod-Giannone M-C, Dallet L, Naudin G, Sahin A, Decossas M, Poussard S and Lambert O. Involvement of caveolin-1 and CD36 in native LDL endocytosis by endothelial cells. Biochimica et Biophysica Acta (BBA)-General Subjects. 2019;1863:830–838.

55. Shvets E, Bitsikas V, Howard G, Hansen CG and Nichols BJ. Dynamic caveolae exclude bulk membrane proteins and are required for sorting of excess glycosphingolipids. Nature communications. 2015;6:6867.

56. Lim JE, Bernatchez P and Nabi IR. Scaffolds and the scaffolding domain: an alternative paradigm for caveolin-1 signaling. Biochemical Society Transactions. 2024:BST20231570.

57. Stamler JS. Redox signaling: nitrosylation and related target interactions of nitric oxide. Cell. 1994;78:931–936.

58. Forrester MT, Foster MW, Benhar M and Stamler JS. Detection of protein S-nitrosylation with the biotin-switch technique. Free Radical Biology and Medicine. 2009;46:119–126.

59. Xue Y, Liu Z, Gao X, Jin C, Wen L, Yao X and Ren J. GPS-SNO: computational prediction of protein S-nitrosylation sites with a modified GPS algorithm. PloS one. 2010;5:e11290.

60. Pepino MY, Kuda O, Samovski D and Abumrad NA. Structure-function of CD36 and importance of fatty acid signal transduction in fat metabolism. Annual review of nutrition. 2014;34:281–303.

61. May SC and Sahoo D. Now in 3D! Novel insights into CD36 structure and function. Ann Blood. 2021;6:1–12.

62. Ottolini M, Daneva Z, Chen YL, Cope EL, Kasetti RB, Zode GS and Sonkusare SK. Mechanisms underlying selective coupling of endothelial Ca2+ signals with eNOS vs. IK/SK channels in systemic and pulmonary arteries. The Journal of physiology. 2020;598:3577–3596.

63. de Wit C, Jahrbeck B, Schäfer C, Bolz S-S and Pohl U. Nitric oxide opposes myogenic pressure responses predominantly in large arterioles in vivo. Hypertension. 1998;31:787–794.

64. Bolz SS, De Wit C and Pohl U. Endothelium-derived hyperpolarizing factor but not NO reduces smooth muscle Ca2+ during acetylcholine-induced dilation of microvessels. British journal of pharmacology. 1999;128:124–134.

65. Garland J and McPherson GA. Evidence that nitric oxide does not mediate the hyperpolarization and relaxation to acetylcholine in the rat small mesenteric artery. British journal of pharmacology. 1992;105:429.

66. Earley S, Pauyo T, Drapp R, Tavares MJ, Liedtke W and Brayden JE. TRPV4-dependent dilation of peripheral resistance arteries influences arterial pressure. American Journal of Physiology-Heart and Circulatory Physiology. 2009;297:H1096–H1102.

67. Razani B, Engelman JA, Wang XB, Schubert W, Zhang XL, Marks CB, Macaluso F, Russell RG, Li M and Pestell RG. Caveolin-1 null mice are viable but show evidence of hyperproliferative and vascular abnormalities. Journal of Biological Chemistry. 2001;276:38121–38138.

68. Cohen AW, Razani B, Wang XB, Combs TP, Williams TM, Scherer PE and Lisanti MP. Caveolin-1-deficient mice show insulin resistance and defective insulin receptor protein expression in adipose tissue. American Journal of Physiology-Cell Physiology. 2003;285:C222–C235.

69. Pojoga LH, Underwood PC, Goodarzi MO, Williams JS, Adler GK, Jeunemaitre X, Hopkins PN, Raby BA, Lasky-Su J and Sun B. Variants of the caveolin-1 gene: a translational investigation linking insulin resistance and hypertension. The Journal of Clinical Endocrinology & Metabolism. 2011;96:E1288–E1292.

70. Ibrahimi A, Bonen A, Blinn WD, Hajri T, Li X, Zhong K, Cameron R and Abumrad NA. Muscle-specific overexpression of FAT/CD36 enhances fatty acid oxidation by contracting muscle, reduces plasma triglycerides and fatty acids, and increases plasma glucose and insulin. Journal of Biological Chemistry. 1999;274:26761–26766.

71. Lohman AW, Straub AC and Johnstone SR. Identification of connexin43 phosphorylation and S-nitrosylation in cultured primary vascular cells. Gap Junction Protocols. 2016:97–111.

72. Thorne RF, Ralston KJ, de Bock CE, Mhaidat NM, Zhang XD, Boyd AW and Burns GF. Palmitoylation of CD36/FAT regulates the rate of its post-transcriptional processing in the endoplasmic reticulum. Biochimica et Biophysica Acta (BBA)-Molecular Cell Research. 2010;1803:1298–1307.

73. Jiang Y, Krantz S, Qin X, Li S, Gunasekara H, Kim Y-M, Zimnicka A, Bae M, Ma K and Toth PT. Caveolin-1 controls mitochondrial damage and ROS production by regulating fission-fusion dynamics and mitophagy. Redox Biology. 2022;52:102304.

74. Schiattarella GG, Altamirano F, Tong D, French KM, Villalobos E, Kim SY, Luo X, Jiang N, May HI and Wang ZV. Nitrosative stress drives heart failure with preserved ejection fraction. Nature. 2019;568:351–356.

75. Opie LH. Metabolic syndrome. Circulation. 2007;115:e32–e35.

76. Denisenko YK, Kytikova OY, Novgorodtseva TP, Antonyuk MV, Gvozdenko TA and Kantur TA. Lipid-induced mechanisms of metabolic syndrome. Journal of Obesity. 2020;2020.

77. Hink U, Li H, Mollnau H, Oelze M, Matheis E, Hartmann M, Skatchkov M, Thaiss F, Stahl RA and Warnholtz A. Mechanisms underlying endothelial dysfunction in diabetes mellitus. Circulation research. 2001;88:e14–e22.

78. Romero N, Radi R, Linares E, Augusto O, Detweiler CD, Mason RP and Denicola A. Reaction of human hemoglobin with peroxynitrite: isomerization to nitrate and secondary formation of protein radicals. Journal of Biological Chemistry. 2003;278:44049–44057.

79. Ottolini M, Hong K, Cope EL, Daneva Z, DeLalio LJ, Sokolowski JD, Marziano C, Nguyen NY, Altschmied J and Haendeler J. Local peroxynitrite impairs endothelial transient receptor potential vanilloid 4 channels and elevates blood pressure in obesity. Circulation. 2020;141:1318–1333.

80. Daneva Z, Marziano C, Ottolini M, Chen Y-L, Baker TM, Kuppusamy M, Zhang A, Ta HQ, Reagan CE and Mihalek AD. Caveolar peroxynitrite formation impairs endothelial TRPV4 channels and elevates pulmonary arterial pressure in pulmonary hypertension. Proceedings of the National Academy of Sciences. 2021;118:e2023130118.

81. Potje SR, Grando MD, Chignalia AZ, Antoniali C and Bendhack LM. Reduced caveolae density in arteries of SHR contributes to endothelial dysfunction and ROS production. Scientific reports. 2019;9:6696.

82. Kang DH, Kim Y, Min S, Lee SY, Chung KY, Baek I-J, Kwon K, Jo H and Kang SW. Blood flow patterns switch VEGFR2 activity through differential S-nitrosylation and S-oxidation. Cell Reports. 2023;42.

83. Toth PP, Patti AM, Giglio RV, Nikolic D, Castellino G, Rizzo M and Banach M. Management of statin intolerance in 2018: still more questions than answers. American Journal of Cardiovascular Drugs. 2018;18:157–173.

84. Cao G, Yang G, Timme TL, Saika T, Truong LD, Satoh T, Goltsov A, Park SH, Men T and Kusaka N. Disruption of the caveolin-1 gene impairs renal calcium reabsorption and leads to hypercalciuria and urolithiasis. The American journal of pathology. 2003;162:1241–1248.

85. DeLalio LJ, Keller AS, Chen J, Boyce AK, Artamonov MV, Askew-Page HR, Keller IV TS, Johnstone SR, Weaver RB and Good ME. Interaction between pannexin 1 and caveolin-1 in smooth muscle can regulate blood pressure. Arteriosclerosis, thrombosis, and vascular biology. 2018;38:2065–2078.

86. Kenwood BM, Weaver JL, Bajwa A, Poon IK, Byrne FL, Murrow BA, Calderone JA, Huang L, Divakaruni AS and Tomsig JL. Identification of a novel mitochondrial uncoupler that does not depolarize the plasma membrane. Molecular metabolism. 2014;3:114–123.

87. Billaud M, Lohman AW, Straub AC, Parpaite T, Johnstone SR and Isakson BE. Characterization of the thoracodorsal artery: morphology and reactivity. Microcirculation. 2012;19:360–372.

88. Wan J, Roth AF, Bailey AO and Davis NG. Palmitoylated proteins: purification and identification. Nature protocols. 2007;2:1573–1584.

